# Regulation of lactose and galactose growth: Insights from a unique metabolic gene cluster in *Candida intermedia*

**DOI:** 10.1101/2023.12.19.572343

**Authors:** Kameshwara V. R. Peri, Le Yuan, Fábio Faria Oliveira, Karl Persson, Hanna D Alalam, Lisbeth Olsson, Johan Larsbrink, Eduard J Kerkhoven, Cecilia Geijer

**Affiliations:** Department of Life Sciences, Chalmers University of Technology, Gothenburg, Sweden; Wallenberg Wood Science Center, Chalmers University of Technology, 412 96, Gothenburg, Sweden; Novo Nordisk Foundation Center for Biosustainability, Technical University of Denmark, DK-2800 Kgs. Lyngby, Denmark; SciLifeLab, Chalmers University of Technology, 41296, Gothenburg, Sweden

**Keywords:** Cheese whey, metabolism, evolution, gene clusters, transcriptional regulation, galactose regulatory system, non-conventional yeast

## Abstract

Lactose assimilation is a relatively rare trait in yeasts, and *Kluyveromyces* yeast species have long served as model organisms for studying lactose metabolism. Meanwhile, the metabolic strategies of most other lactose-assimilating yeasts remain unknown. In this work, we have elucidated the genetic determinants of the superior lactose-growing yeast *Candida intermedia*. Through genomic and transcriptomic analyses and deletion mutant phenotyping, we identified three interdependent gene clusters responsible for the metabolism of lactose and its hydrolysis product galactose: the conserved *LAC* cluster (*LAC12, LAC4*) for lactose uptake and hydrolysis, the conserved *GAL* cluster (*GAL1, GAL7, GAL10*) for galactose catabolism, and a unique “*GALLAC”* cluster. This novel *GALLAC* cluster, which has evolved through gene duplication and divergence, proved indispensable for *C. intermedia’s* growth on lactose and galactose. The cluster contains the transcriptional activator gene *LAC9*, second copies of *GAL1* and *GAL10* and the *XYL1* gene encoding an aldose reductase involved in carbon overflow metabolism. Notably, the regulatory network in *C. intermedia*, governed by Lac9 and Gal1 from the *GALLAC* cluster, differs significantly from the (ga)lactose regulons in *Saccharomyces cerevisiae*, *Kluyveromyces lactis* and *Candida albicans*. Moreover, although lactose and galactose metabolism are closely linked in *C. intermedia*, our results also point to important regulatory differences. This study paves the way to a better understanding of lactose and galactose metabolism in *C. intermedia* and provides new evolutionary insights into yeast metabolic pathways and regulatory networks. In extension, the results will facilitate future development and use of *C. intermedia* as a cell-factory for conversion of lactose-rich whey into value-added products.

## Introduction

Assimilation of lactose is a rather uncommon characteristic among microorganisms, including yeasts. Growth screening of 332 genome-sequenced yeasts from the Ascomycota phylum showed that only 24 (<10%) could grow on lactose, and these lactose-utilizers are scattered throughout the phylogenetic tree^1^. While lactose increased in abundance on earth with the domestication of lactating mammals about 10,000 years ago^2^, ascomycetous yeast clades formed already millions of years ago^1^, suggesting that lactose metabolism may have evolved several times throughout yeast evolution. Whereas ‘dairy yeast’ from the *Kluyveromyces* genus, including *K. lacti*s and *K. marxianus*, have been carefully characterized^3–6^, other lactose-metabolizing yeast species remain largely understudied. Elucidating the mechanisms behind their lactose metabolism can help to shed light on how eukaryotic metabolic pathways and the associated regulatory networks have evolved. Moreover, it can enable the development of new yeast species as cell factories for conversion of lactose in the abundant industrial side stream cheese whey into a range of different products^7^.

Lactose is a disaccharide composed of D-glucose and D-galactose connected through a β-1,4-glycosidic linkage. Its assimilation starts with the hydrolysis of lactose into its monosaccharides through the action of a lactase – normally an enzyme with β-galactosidase activity. Several different enzyme families encode lactases, which can be found intracellularly or extracellularly. In *Kluyveromyces* yeasts, lactose is transported across the plasma membrane by a *LAC12*-encoded lactose permease and is subsequently hydrolyzed intracellularly by a *LAC4*-encoded *β*-galactosidase^6^. In contrast, the yeast *Moesziomyces aphidis* and *M. antarcticus* seem to show *β*-galactosidase activity both intra and extracellularly, whereafter glucose and galactose are imported into the cell^8^. For most other lactose-growing yeast, comparative genomics and growth characterization are still needed to determine their lactose uptake and hydrolysis mechanisms.

In *Kluyveromyces* (and likely most other lactose-assimilating yeasts), lactose-derived glucose and galactose moieties are further catabolized through glycolysis and the Leloir pathway, respectively. The Leloir pathway is carried out by Gal1, Gal7 and Gal10, and starts by conversion of β-D-galactose into α-D-galactose by the mutarotase domain of Gal10 (aldose-1-epimerase). Gal1 (galactokinase) then phosphorylates α-D-galactose into α-D-galactose-1-phosphate, whereafter Gal7 (galactose-1-phosphate uridylyl transferase) transfers uridine diphosphate (UDP) from UDP-α-D-glucose-1-phosphate to α-D-galactose-1-phosphate^9^. The epimerase (UDP-galactose-4-epimerase) domain of Gal10 catalyzes the final step, where UDP-α-D-galactose-1-phosphate is converted to UDP-α-D-glucose-1-phosphate^10–12^. In parallel to the Leloir pathway, some filamentous fungi such as *Trichoderma reesei* and *Aspergillus nidulans* have an alternative galactose catabolic pathway called the oxidoreductive pathway, where galactose is first converted into galactitol through the action of an aldose reductase^11,12^. Also a third galactose catabolic pathway, the DeLey-Doudoroff pathway, has been described to some detail^12^. To the best of our knowledge, (ga)lactose-growing yeasts described to date exclusively use the Leloir pathway, although some reports on galactose-to-galactitol conversion in *Rhodosporidium toruloides* and *Metschnikowia pulcherrima* exist^13–15^. Moreover, 12 out of 332 ascomycetous yeasts have been shown to grow on galactitol^1^, indicating that they might possess an oxidoreductive pathway to catabolize this carbon source.

Comparative genomic studies have revealed that the *GAL1, 7* and *10* genes are often found located together in a “*GAL* cluster” in the genomes of yeast and filamentous fungi^16^, and also the *LAC4* and *LAC12* genes form a “*LAC* cluster” in for example *K. marxianus* and *K. lactis*^6,16^. Such metabolic gene clusters, identified both in filamentous fungi and yeasts, are particularly prevalent for pathways involved in sugar and nutrient acquisition, synthesis of vitamins and secondary metabolites^17^. Some clusters, including the *GAL* cluster, are conserved over a wide range of species whereas other clusters appear unique to one or a few species^16,18,19^. Like bacterial operons, the eukaryotic cluster genes are co-regulated in response to environmental changes, and clusters sometimes even encode their own transcriptional activators^17^. Clustering of genes under a common control mechanism allows the microorganism to rapidly adapt to environmental cues, which can be advantageous to avoid deleterious recombination events and high concentrations of local protein products. For example, co-regulation of the *GAL* genes is necessary to avoid accumulation of the toxic intermediate galactose-1-phosphate in the Leloir pathway^16,20^. Gene clusters can also propagate together by horizontal transfers to other species, which is less likely to occur for non-clustered genes^21^. In fact, selective pressures in lactose-rich environments in dairy farms led to the formation of an efficient lactose utilization system by rearrangement and horizontal gene transfer (HGT) of the *LAC* cluster genes in *Kluyveromyces* dairy strains^6^.

Regulation of galactose metabolism (and lactose where applicable) has been carefully characterized in yeasts such as *S. cerevisiae, K. lactis* and *Candida albicans* ^22–25^, displaying both similarities and differences among species. In *S. cerevisiae*, three regulatory proteins (*Sc*Gal4, *Sc*Gal80, *Sc*Gal3) are responsible for galactose regulation. In the absence of galactose, the transcriptional activation domain of *Sc*Gal4 is bound to the inhibitor *Sc*Gal80. In the presence of galactose, *Sc*Gal3 relieves *Sc*Gal4 from *Sc*Gal80 in a galactose- and ATP-dependent manner, resulting in the induction of the *GAL* structural genes. Like for *S. cerevisiae*, *K. lactis GAL* regulatory system relies on relieving *Kl*Lac9 (ortholog of *Sc*Gal4) from Gal80 inhibition. However, *K. lactis* lacks Gal3 and instead uses a bifunctional galactokinase *Kl*Gal1 to induce both galactose and lactose genes^26^. There are four *Kl*Lac9 binding sites in the *LAC* cluster gene promoters, which indicate the tight coregulation of lactose and galactose metabolism in this yeast^27^. Similar to *K. lactis*, *C. albicans* lacks Gal3 but possesses a Gal1 with both enzymatic and regulatory functions, but in this yeast the *GAL* gene expression is controlled by transcription factors Rtg1/Rtg3^28^ and/or *Ca*Rep1/*Ca*Cga1^29^ rather than *Ca*Gal4, which instead is responsible for expression of genes involved in glucose metabolism^22^. Such transcriptional rewiring is common among yeasts, which calls for coupling of comparative genomics with detailed mutant phenotyping and transcriptional analysis to decipher how regulation occurs in individual species.

While (ga)lactose metabolism in *S. cerevisiae* and *K. lactis* has long served as a model system for understanding the function, evolution and regulation of eukaryotic metabolic pathways, the corresponding knowledge regarding non-conventional yeasts is scarce. One such non-conventional yeast is *Candida intermedia*, a haploid yeast belonging to the *Metschnikowia* family in the CUG-Ser1 clade, which can grow on a wide range of different carbon sources^1^. *C. intermedia* has previously received attention as a fast-growing yeast on xylose. The xylose transporters and xylose reductases responsible for *C. intermedia’s* xylose-fermentative capacity have been characterized in several studies^30–35^. *C. intermedia* is one of very few yeasts in the *Metschnikowia* family that can grow on lactose^1^, and it has been used for cheese whey bioremediation in the past^36^. Our previous works on characterizing the in-house isolated *C. intermedia* strain CBS 141442 in terms of genomics, transcriptomics and physiology^33,37,38^ and the development of a genome editing toolbox for this species^39^ provide a stable platform for exploration of the genetic determinants of lactose metabolism in this yeast.

In this study, we show that *C. intermedia* possesses a unique ‘*GALLAC*’ cluster, in addition to the conserved *GAL* and *LAC* clusters, that is essential for growth on lactose and highly important for growth on galactose. Characterization of the individual genes within *GALLAC* cluster revealed differentiation in their functionality, enabling the yeast to regulate the expression of galactose and lactose genes differently. This cluster represents a new, interesting example of metabolic network rewiring in yeast, and likely helps to explain how *C. intermedia* has evolved into an efficient lactose-assimilating yeast.

## Results

### C. intermedia is among the top five lactose-growers out of 332 sequenced ascomycetous yeasts

As a start, we wanted to assess the capacity of *C. intermedia* to grow on lactose compared to other yeasts. We cultured 24 of the 332 ascomycetous species that have scored positive for lactose growth^1^, as well as *C. intermedia* strains CBS 572 (type strain), CBS 141442 and PYCC 4715 (previously characterized for utilization of xylose)^1,34^. The yeast species displayed different growth patterns in lag phase, doubling time and final biomass (Figure 1, Figure S 1). When ranked based on lowest doubling time, *K. lactis* and *K. marxianus* were the fastest growers on lactose, closely followed by *C. intermedia* strains PYCC 4715 and CBS 141442, *Debaryomyces subglobulus* and *Blastobotrys muscicola* (Figure 1, Figure S 1). Other species such as *Kluyveromyces aestuarii*, *Millerozyma acaciae* and *Lipomyces mesembris* showed poor or no growth under the conditions tested while others had very long lag phases. Thus, under the assessed conditions, our results establish *Candida intermedia* as one of the top five fastest lactose-growing species within this subset of ascomycetous yeasts^1^.

**Figure 1:**
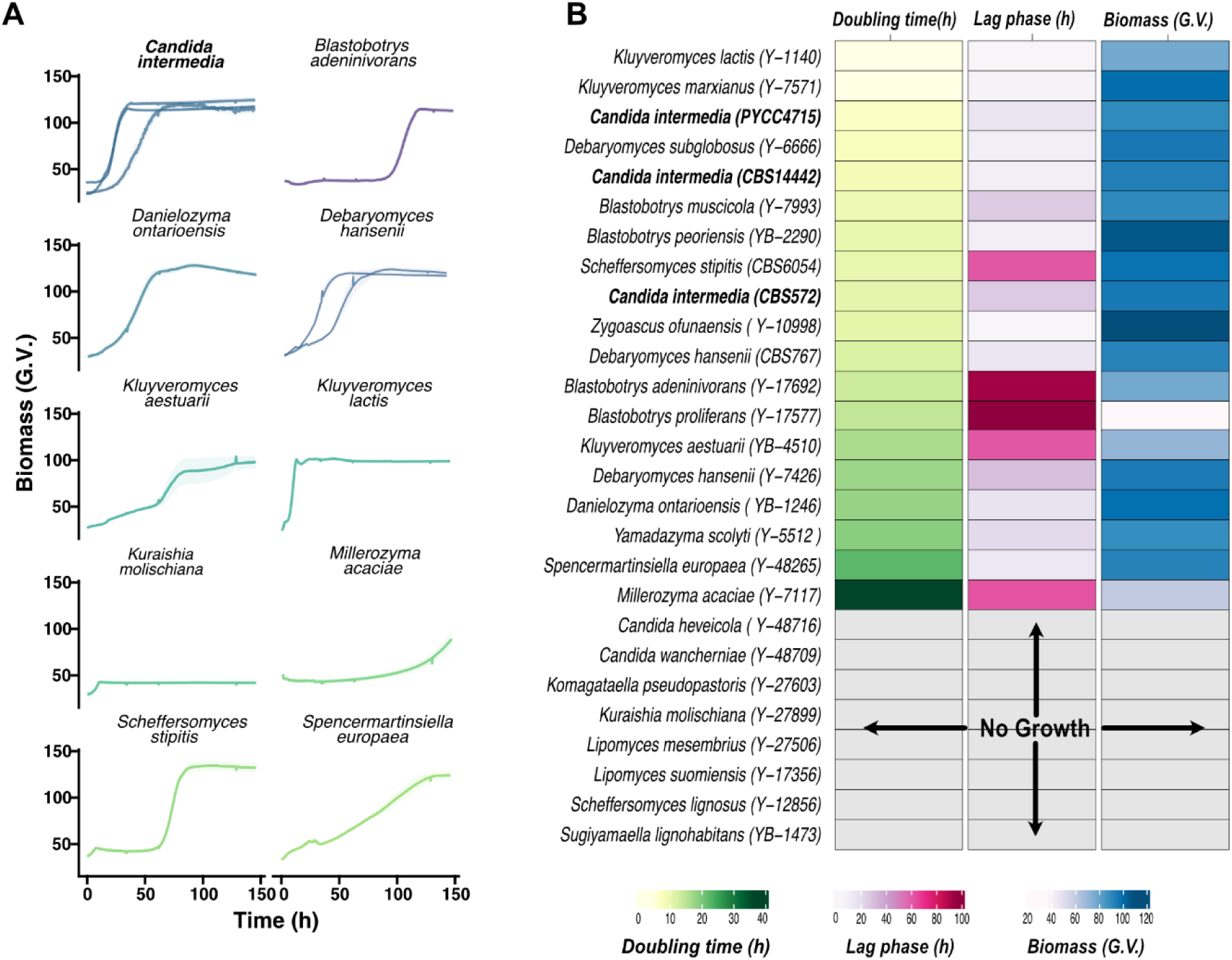
*Candida intermedia* is one of the top five fastest lactose-growing yeast species. A) Representative growth profiles of 10/24 lactose-yeast species including three different *C. intermedia* strains. The graphs depict data procured from GrowthProfiler in 96-well format, represented as mean ± standard deviation (shaded region) for biological triplicates. On y-axis final biomass is depicted in green values (G.V. - corresponding to growth based on pixel counts, as determined by a GrowthProfiler instrument) and is plotted against time (h) on x-axis. B) Heat map showing doubling time (h), lag phase duration (h) and final biomass (green values – G.V.) measured for all the tested strains in minimal media containing lactose as the sole carbon source and plotted as an average of three biological replicates. Strains are ranked based on their doubling time, from low to high.

### Genomic and transcriptomic analysis identify three gene clusters involved in lactose and galactose assimilation

To identify the genetic determinants for lactose metabolism in *C. intermedia* CBS 141442, we searched the genome for orthologs of known genes involved in the uptake and conversion of lactose, and its tightly coupled hydrolysis-product, galactose. We found several genes encoding expected transcription factor orthologs including *LAC9, GAL4, RTG1, RTG3, REP1* and *CGA1* that have been associated with lactose and galactose metabolism in *K. lactis*^40^, *S. cerevisiae*^41^ and *C. albicans*^28,29^. In accordance with previous reports for yeasts belonging to the genus *Candida*^10^, we did not find orthologous of *GAL80*, strongly suggesting that *C. intermedia* does not possess the Gal3-Gal80-Gal4 regulon.

Moreover, genome of *C. intermedia* contains the conserved *GAL* cluster including *GAL1, 7, 10* genes as well as an *ORF-X* gene encoding a putative glucose-4,6-dehydratase similar to *GAL* clusters in *Candida/Schizosaccharomyces* strains^10,16^ (Figure 2). We also identified the conserved *LAC* cluster containing the *β*-galactosidase gene *LAC4* and lactose permease gene *LAC12*^3,4^, which correlates well with *C. intermedia* predominantly displaying intracellular *β*-galactosidase activity (data not shown)^6^. To our surprise, *C. intermedia* also possesses a third cluster, hereafter referred to as the *GALLAC* cluster, containing a putative transcriptional regulator gene *LAC9 (LAC9_2)* next to a second copy of the *GAL1* gene (*GAL1_2*), followed by one of the three xylose/aldose reductase genes (*XYL1_2*) previously characterized in *C. intermedia*^37^ and lastly, a second copy of *GAL10* (*GAL10_2*). Interestingly, the *GAL10_2* gene is shorter than *GAL10* in the *GAL* cluster and seems to encode only the epimerase domain, similar to *GAL10* orthologs in *Schizosaccharomyces* species and filamentous fungi^10^.

**Figure 2:**
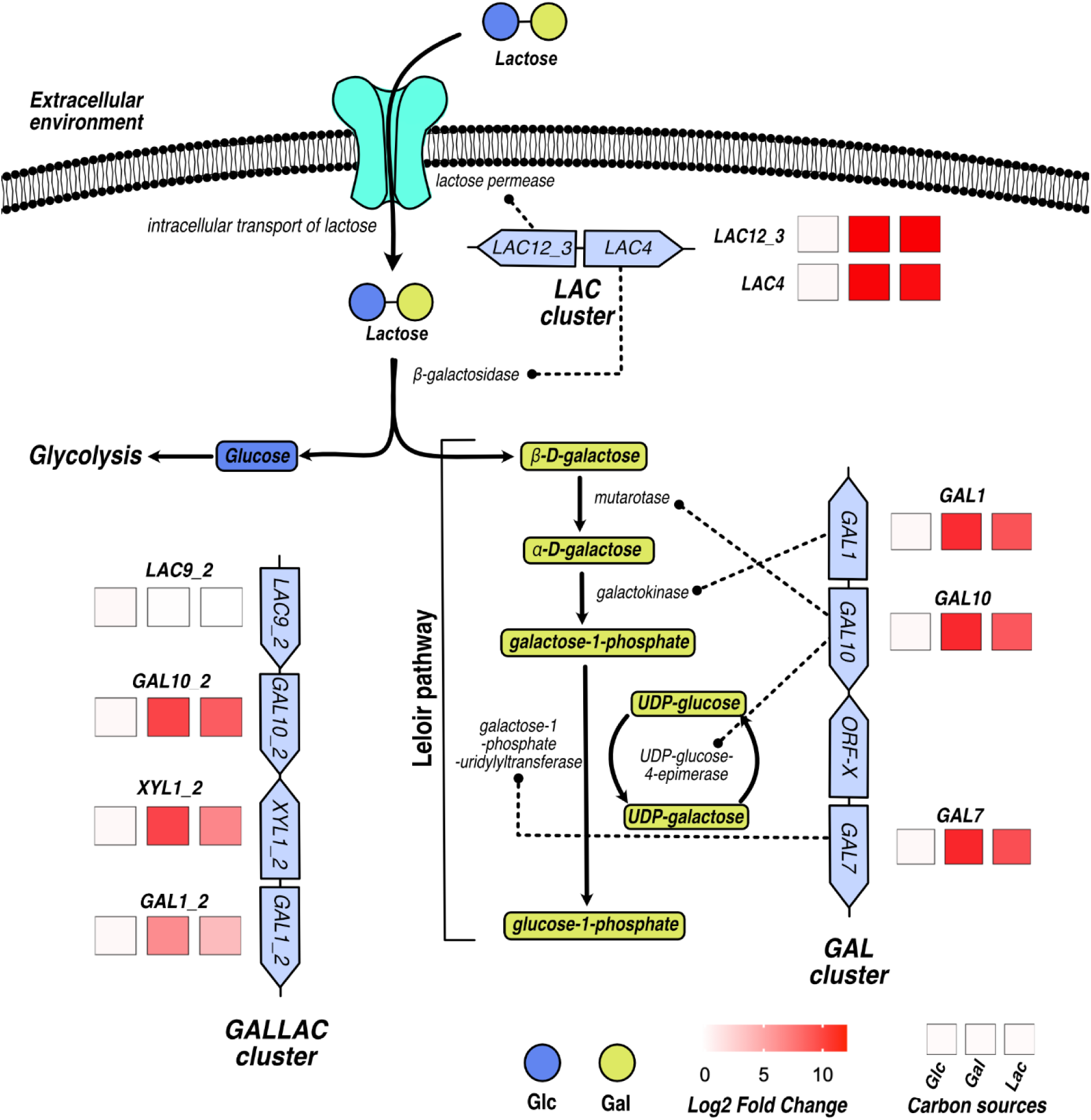
Genomic and transcriptomic analyses identified three gene clusters involved in lactose and galactose assimilation: A schematic representation of lactose and galactose metabolic pathways and results of RNAseq data analysis showing expression of different genes (as present in clusters) upregulated in galactose or lactose compared to glucose. Lactose uptake and transport into the cell is enabled by *LAC12_3* encoded lactose permease followed by hydrolysis to glucose (blue circle: Glc) and galactose (yellow circle: Gal) enabled by *LAC4* encoded β-galactosidase enzyme. Glucose is further metabolized via glycolysis. Galactose is metabolized via the Leloir pathway, encoded by three clustered genes, *GAL1* (galactokinase), *GAL7* (galactose-1-phosphate-uridylyltransferase) and *GAL10* (mutarotase and UDP-glucose-4-epimerase). The enzymatic functions for the genes are depicted by dotted lines based on genome sequence data for *C. intermedia* CBS141442. Legend shows Log2 fold change with carbon sources tested represented as Glc for 2% glucose, Gal for 2% galactose and Lac for 2% lactose containing media. Gene expression log fold change is normalized with glucose as control.

Next, we performed transcriptome analysis using RNA-sequencing (RNA-seq) technology on the CBS 141442 strain cultivated in media containing 2% of either lactose, galactose, or glucose (Figure 2). All genes in the *LAC* and *GAL* clusters were among the highest upregulated genes in both galactose and lactose as compared to glucose conditions. Also, the genes in the *GALLAC* cluster were highly upregulated on both of these carbon sources with respect to glucose, with the exception of the constitutively expressed *LAC9_2* gene (Figure S 2), indicating that the novel cluster might play an important role in galactose and lactose metabolism in this non-conventional yeast.

### The GALLAC cluster is essential for growth on lactose and unique to C. intermedia

To decipher the importance of the three clusters for (ga)lactose metabolism in *C. intermedia*, we deleted the clusters one by one using the split-marker technique previously developed for this yeast^39^. The cluster deletion mutants (*lacΔ*, *galΔ* and *gallacΔ*) grew almost as well as the wild-type strain (WT) in minimal media containing glucose (Figure 3A). As expected, *galΔ* failed to grow on galactose, which can be attributed to the complete shut-down of the Leloir pathway, whereas the *lacΔ* grew like WT. Interestingly, no growth was observed for the *gallacΔ* in galactose during the first 90 h, whereafter it slowly started to grow (Figure 3B). With lactose as carbon source, both *lacΔ* and *gallacΔ* completely failed to grow, whereas *galΔ* started to grow slowly after approx. 100 h (Figure 3C). Thus, our results show that the *GALLAC* cluster is essential for growth on lactose and highly important for growth on galactose.

**Figure 3:**
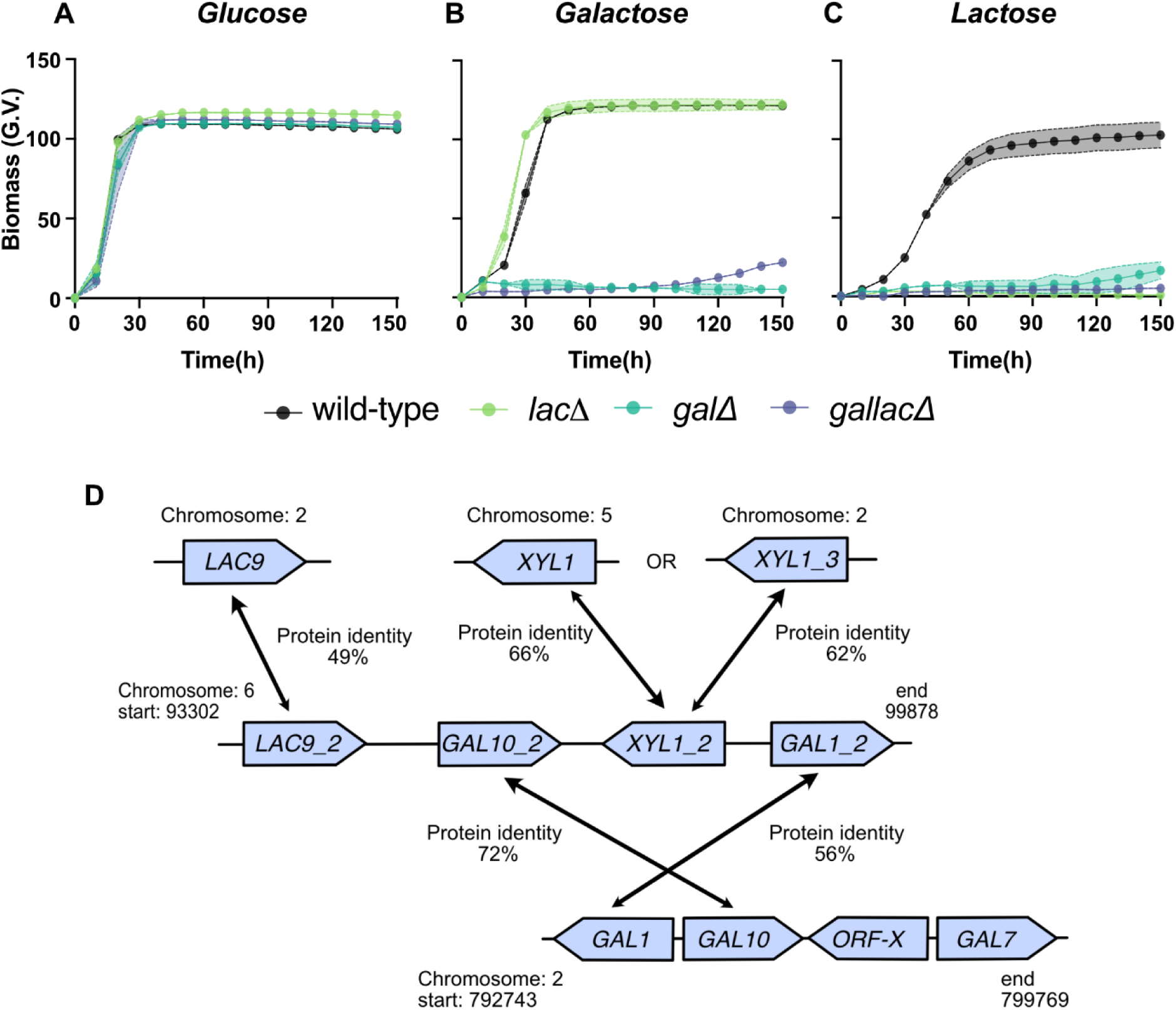
The *GALLAC* cluster is essential for growth on lactose and galactose and is unique to *C. intermedia*: Cluster deletion mutants of *C. intermedia* were characterized by growth on glucose (A), galactose (B) lactose (C) in growth profiler. Legend shows the wild-type strain (black), *LAC* cluster mutant (light green), *GAL* cluster deletion mutant (dark green) and *GALLAC* cluster deletion mutant (purple), depicted in the graph with biomass as green values (G.V. - corresponding to growth based on pixel counts, as determined by a GrowthProfiler instrument) on the y-axis against time(h) on the x-axis. Data are represented as mean ± standard deviation (shaded region) for biological triplicates indicated by colors: wild type – black, lac cluster mutant – light green, gal cluster mutant – dark green and gallac cluster mutant – purple. D) Graphical representation of genomic location of cluster and individual genes which are paralogs to *GALLAC* gene cluster and their protein identity as per comparative genomics analysis. Arrows depict assumed duplication events which are still unclear.

To the best of our knowledge, the existence of a *GALLAC*-like cluster and its interdependence with the *GAL* and *LAC* clusters has never previously been reported. This, along with the severe growth defects of *gallacΔ,* encouraged us to determine the origin and prevalence of the cluster in other yeasts. First, we performed a comparative genomic analysis among the dataset of 332 genome-sequenced ascomycetous yeasts^1^. Although *GAL1* and *GAL10* were found clustered together as parts of the conserved *GAL* clusters in 150/332 species^16^, *C. intermedia* was the only species where these genes also clustered with *LAC9* and *XYL1* genes (Figure 3D). Next, to decipher the evolutionary events that led to the formation of the *GALLAC* cluster, we generated phylogenetic trees for each individual gene product of the cluster. Our analysis revealed that although the amino acid identities between the paralogs in *C. intermedia* are relatively low (56% for Gal1 and Gal1_2, 72% for Gal110 and Gal10_2, 49% for Lac9_2 and Lac9 and 66% and 62% for Xyl1_2 compared to Xyl1 and Xyl1_3, respectively), the identities between the paralogs are still higher than for most orthologs in other species (Figure 3B, Figure S 3–6). Combined, these results strongly suggest that the unique *GALLAC* cluster has evolved within *C. intermedia* through gene duplication and divergence.

### Deletion of individual genes in the GAL and GALLAC clusters reveals importance of Lac9_2 and Gal1_2 for (ga)lactose metabolism

To elucidate the physiological function of genes situated in the *GALLAC* cluster and to better understand the interdependence between the clusters, we deleted individual genes in both the *GALLAC* and *GAL* clusters. The mutant phenotypes were compared with WT and complete cluster deletions regarding growth, consumption of sugars and production of metabolites in defined media containing either 2% galactose or lactose.

With galactose as carbon source, deletion of *LAC9_2* located in the *GALLAC* cluster resulted in an extended lag phase accompanied by galactitol production, indicating that this putative transcription factor is involved in regulation of galactose metabolism. However, deletion of the other genes in the *GALLAC* cluster did not result in severe growth defects (Figure S 7). For mutants deleted of individual *GAL* cluster genes, we saw the expected severe growth defects for *gal1Δ, gal7Δ* and *gal10Δ* (Figure 4A). However, *gal10Δ* repeatedly displayed some growth after a very long lag phase of approx. 250h (Figure S 8), which could suggest that Gal10_2 from the *GALLAC* cluster can partly complement the deletion of *GAL10* from the *GAL* cluster.

**Figure 4:**
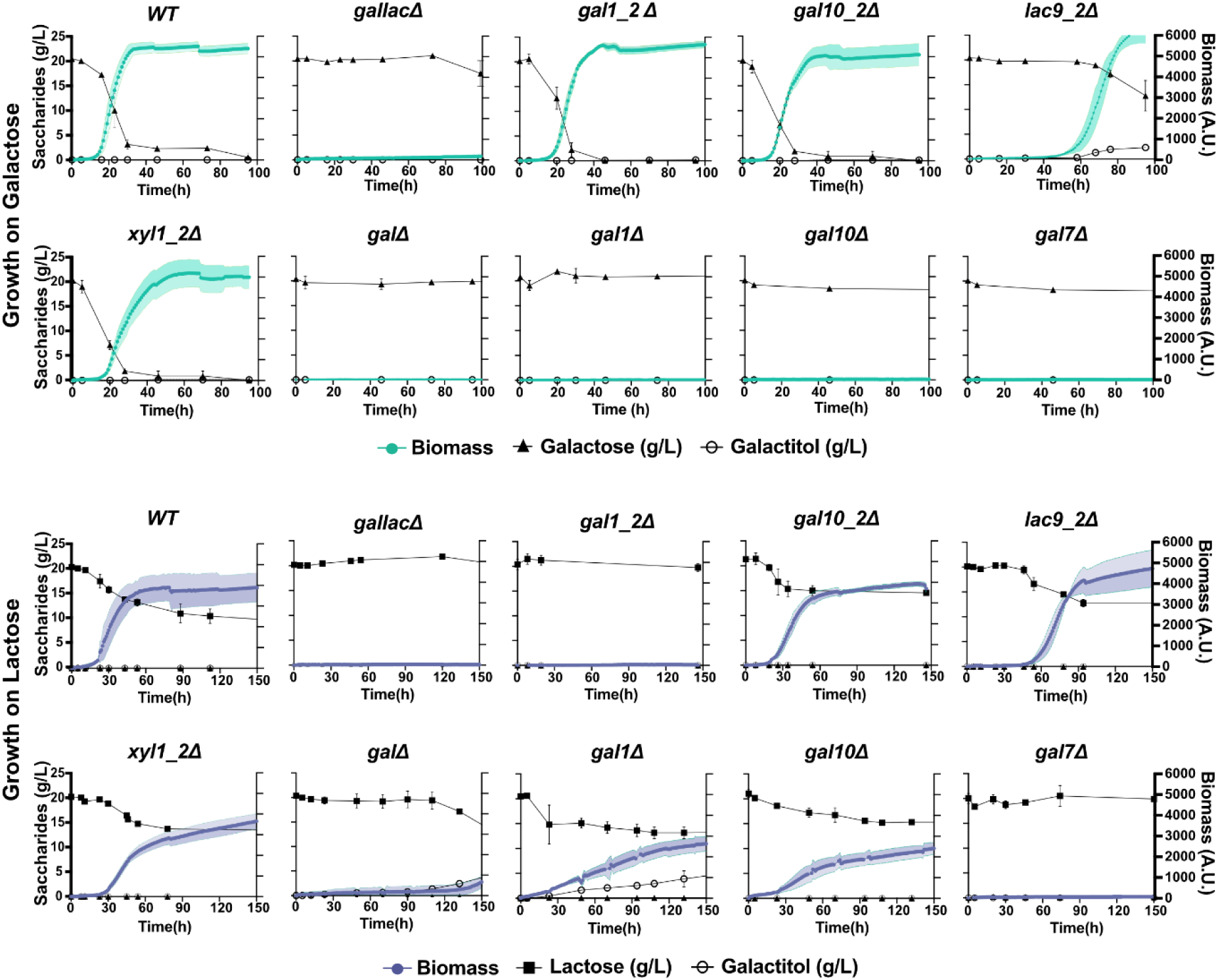
Deletion of individual genes in the *GAL* and *GALLAC* clusters reveals importance of Lac9 and Gal1_2 for (ga)lactose metabolism: Growth and metabolite profiles for deletion mutants of individual genes in the *GAL* and *GALLAC* cluster of *C. intermedia*, in both galactose (top two rows) and lactose (bottom two rows) containing media. Graphs represent biomass (filled circle; gal – dark green; lac-purple) on the right y-axis, consumption of respective sugars (filled triangle for galactose in g/L or filled square for lactose in g/L) and metabolite production (open circle for galactitol in g/L) on the left y-axis (depicted by saccharides (g/L), plotted against time (h) on x-axis. Data are represented as mean ± standard deviation (shaded region for biomass and bars for sugars and metabolites) for biological triplicates.

With lactose as carbon source, *lac9_2* displayed a delay in the onset of growth as was observed for galactose while *gal10_2Δ* and *xyl1_2Δ* grew like WT. However, in contrast to the galactose case, deletion of *GAL1_2* abolished growth and resembled the deletion of the whole *GALLAC* cluster, indicating an important function for this protein in lactose metabolism and a clear phenotypic difference between the two carbon sources. On the contrary, deletion of *GAL1* from the *GAL* cluster did not fully abolish growth on lactose, but growth was slower and accompanied with accumulation of galactitol (73% of theoretical yield), suggesting that most of the lactose-derived galactose is catabolized through the action of an aldose reductase (such as Xyl1_2), rather than through the putative galactokinase Gal1_2 in this mutant. Also, *gal10Δ* grew slowly but with no measurable accumulation of galactose or galactitol, again showing that the *GAL10_2* in the *GALLAC* cluster can partly complement this deletion. Deletion of the only copy of *GAL7* gene encoding for galactose-1-phosphate uridylyltransferase resulted in complete growth inhibition on lactose (as for galactose), and we speculate that the severe growth phenotype is due to the accumulation of toxic intermediate galactose-1-phosphate as seen in *S. cerevisiae* in previous studies ^20^.

As no single deletion resembled the growth defect seen for *gallacΔ* on galactose, we hypothesized that two or more genes must be deleted for the same phenotype to appear. We therefore deleted both *LAC9_2* and *GAL1_2*, which resulted in a growth defect strikingly similar to that of the complete *GALLAC* cluster mutant (Figure S 9). Overall, we can conclude that *Lac9_2* and *gal1_2* have important functions during galactose and lactose growth, although there seem to be significant differences between the two carbon sources.

### Lac9 binding motifs are found in promoters in the GALLAC cluster but not in the GAL and LAC clusters

To better understand the putative role of Lac9_2 as a transcriptional regulator, we performed Multiple Em for Motif Elicitation^42^ (MEME; Version 5.5.43) analysis to identify conserved transcription factor binding motifs in gene promoters in the three clusters. The analysis revealed Lac9 (Gal4) binding motifs (p-value = 8.66×10^-3^) in the promoters of *GAL1_2*, *XYL1_2* and *GAL10_2* in the *GALLAC* cluster, but not in the promoters in the *GAL* and *LAC* clusters (Figure 5A, Figure S 10). These results confirm the bioinformatic analysis of the 332 ascomycetous yeast recently published, showing that *C. intermedia* and many other CTG clade yeasts lack Lac9/Gal4 binding sites in their *GAL* clusters^16^. Although additional analysis would be needed to better understand the transcriptional regulation exerted by Lac9_2, it is likely that it directly binds the promoters of genes within the *GALLAC* cluster.

**Figure 5:**
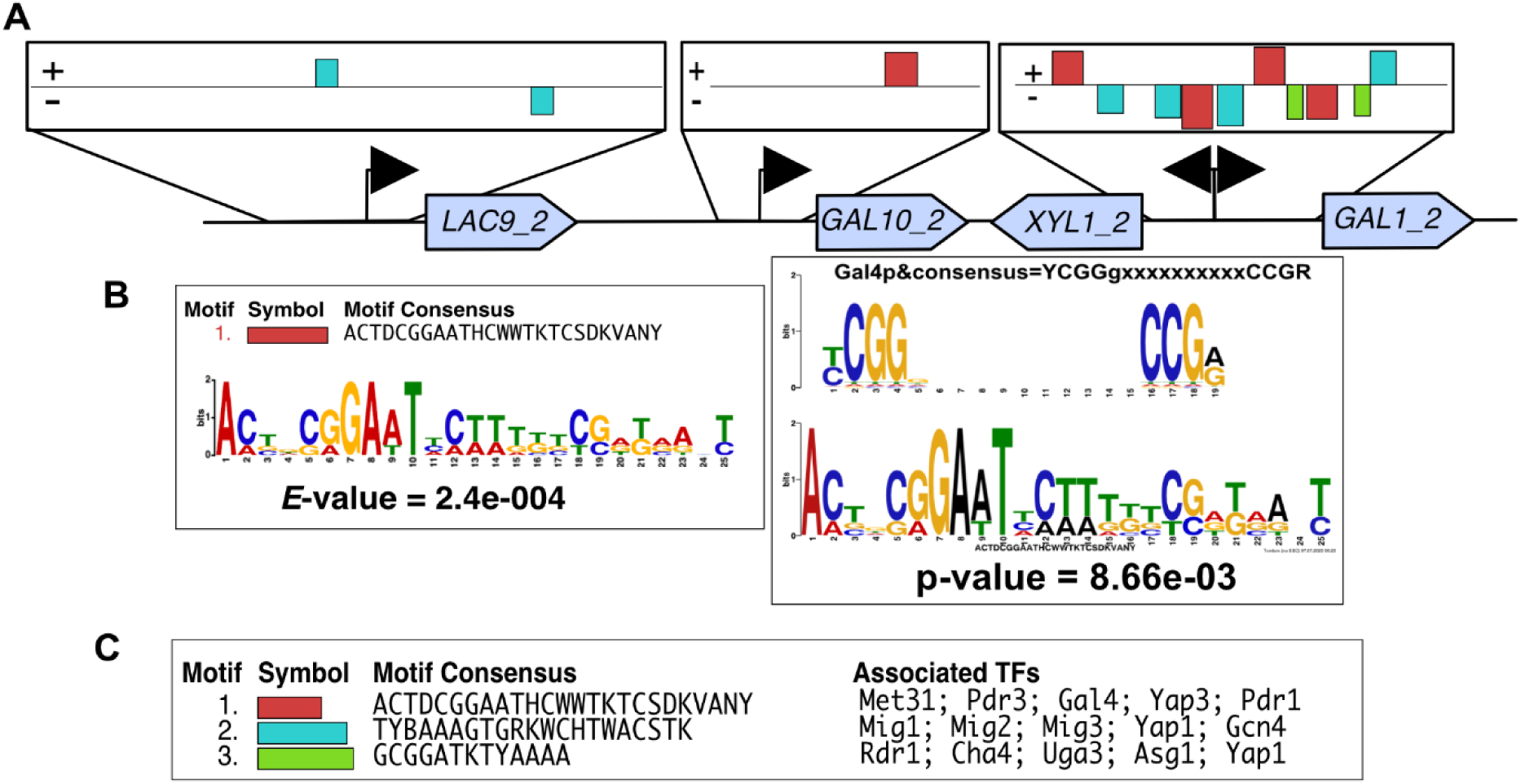
Lac9 binding motifs are found in promoters in the *GALLAC* cluster: A) Graphical representation of results of transcription binding motif analysis for promoters of individual genes of the *GALLAC* cluster, using MEME (version 5.5.43). *GALLAC* gene cluster with the location of three statistically significant promoter binding motifs found in the promoters of the cluster genes. B) Motif consensus of the binding motif with the lowest E-value score of the overall match of the motif in the input sequence. Depiction of the Gal4p consensus sequence and its associated p-value. C) List of three (statistically significant) motifs found in the promoters of *GALLAC* cluster genes and the transcription factors associated to these motifs derived from Yeastract database using TomTom.

Besides *LAC9_2* in the *GALLAC* cluster, our comparative genomics analysis also identified a second, non-clustered *LAC9* gene (Figure 3) as well as *GAL4* gene. All three proteins have predicted Gal4-like DNA-binding domains, but they differ substantially in protein sequence identity (45% for Lac9_2 and Lac9, and 18% and 19% for Gal4 compared to Lac9 and Lac9_2, respectively). As deletion mutants of *LAC9* and *GAL4* did not display growth defects on lactose or galactose (Figure S 11), we conclude that they are not important transcriptional regulators for (ga)lactose metabolism in *C. intermedia*.

### Gal1_2 is required for the induction of LAC cluster genes in C. intermedia

Our deletion mutant phenotyping results suggest that Gal1 and Gal1_2 have at least partly different physiological functions in *C. intermedia* (Figure 4). As both genes are highly upregulated on both galactose and lactose in the WT strain (Figure 2), we speculated that they must differ in their activities as galactokinases or regulators. To this end, we expressed both proteins in *S. cerevisiae* BY4741 *gal1Δ*, which successfully rescued the mutant’s growth defect on galactose (Figure 6A). This experiment demonstrates that both proteins have galactokinase activity, at least when expressed in *S. cerevisiae*. We also compared the predicted structures of Gal1 and Gal1_2 using Alphafold2^43,44^, observing that even though the amino acid sequence identity between the two proteins is as low as 56%, the protein structures are very similar to each other (rmsd 0.490 Å; Figure 6B) as well as to the experimentally solved structure of *Sc*Gal1^42^ (rmsd 0.778 and 0.758 Å for Gal1 and Gal1_2, respectively). Additionally, we observed that the amino acids interacting with galactose in *Sc*Gal1 (PDB ID: 2aj4) are identical to those in the *Ci*Gal1 proteins, apart from Asn213 in *Sc*Gal1 (Asn205 in *Ci*Gal1), which interacts with the O2 hydroxyl group, which in *Ci*Gal1_2 is instead a serine residue (Ser199). The active site clefts of all enzymes are only big enough to accommodate monosaccharides like galactose. Thus, it is highly unlikely that they bind to other, larger substrates such as lactose (Figure 6B). In *S. cerevisiae*, the regulator *Sc*Gal3 is similar in structure to the galactokinase *Sc*Gal1 but has lost its galactokinase activity due to an addition of two extra amino acids (Ser-Ala dipeptide)^45^. However, no such structural changes were seen for *Ci*Gal1 or *Ci*Gal1_2 that could help us to predict regulatory functions.

**Figure 6:**
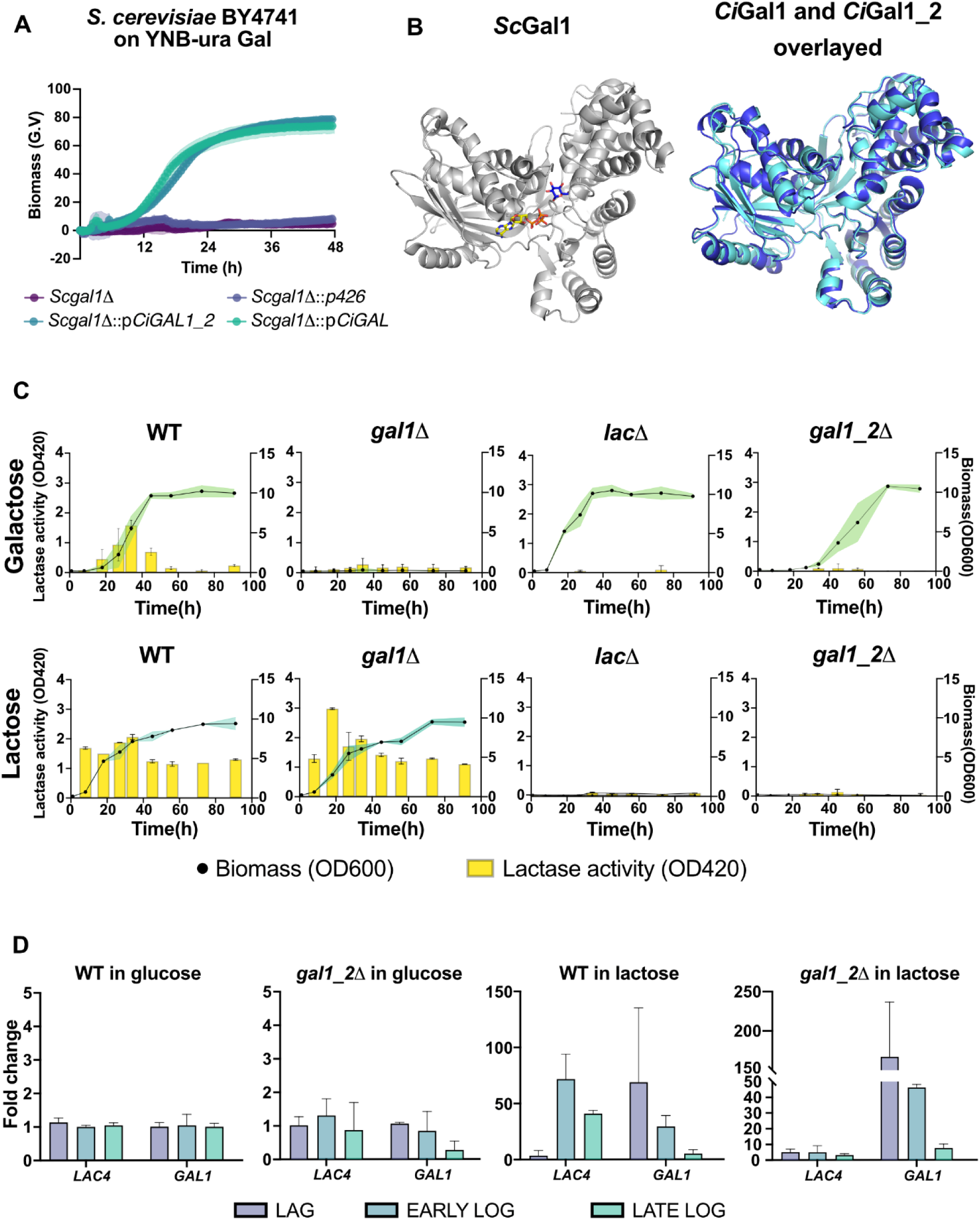
Characterization of *C. intermedia’s* Gal1 and Gal1_2 proteins reveals important functional differences: A) Results of complementation of codon optimized *CiGAL1* and *CiGAL1_2* by heterologous expression in *S. cerevisiae* (BY4741) *gal1Δ* mutant. Growth profiles are depicted for *Scgal1Δ* (dark purple), *Scgal1Δ* with plasmid p426 containing URA marker (light purple), *Scgal1Δ* with *pCiGAL1_2* containing *URA3* marker with codon-optimized *CiGAL1_2*(dark green) and *Scgal1Δ* with *pCiGAL1* containing *URA3* marker with codon-optimized *CiGAL1*). Time (in hours) on x-axis is plotted against biomass (green values – G.V.) on y-axis. Data are represented as mean ± standard deviation (shaded region for biomass) for biological triplicates. B) Structure of ScGal1 (grey) in complex with AMPPNP and α-galactose next to the superimposed Alphafold2-predicted structures of Gal1 (cyan) and Gal1_2 (blue) in the same orientation, showing their high structural similarity C) β-galactosidase assay on galactose- and lactose-grown cultures of wild type, *lacΔ, gal1Δ* and *gal1_2Δ* strains of *C. intermedia*. Graphs show lactase activity (OD420) plotted on left y-axis against time (in hours) on x-axis and biomass (OD_600_) plotted on right y-axis. D) Quantitative PCR results for *LAC4* and *GAL1* gene expression in *C. intermedia* wild-type and *gal1_2Δ* grown in glucose or lactose. Samples were taken during different growth phases (On glucose, Lag = 5h, Early log = 10h and late log = 20h and on lactose, Lag = 5h, early log = 24h, late log = 44h). Data are represented as mean ± standard deviation (error bars) for biological and technical triplicates.

**Figure 7:**
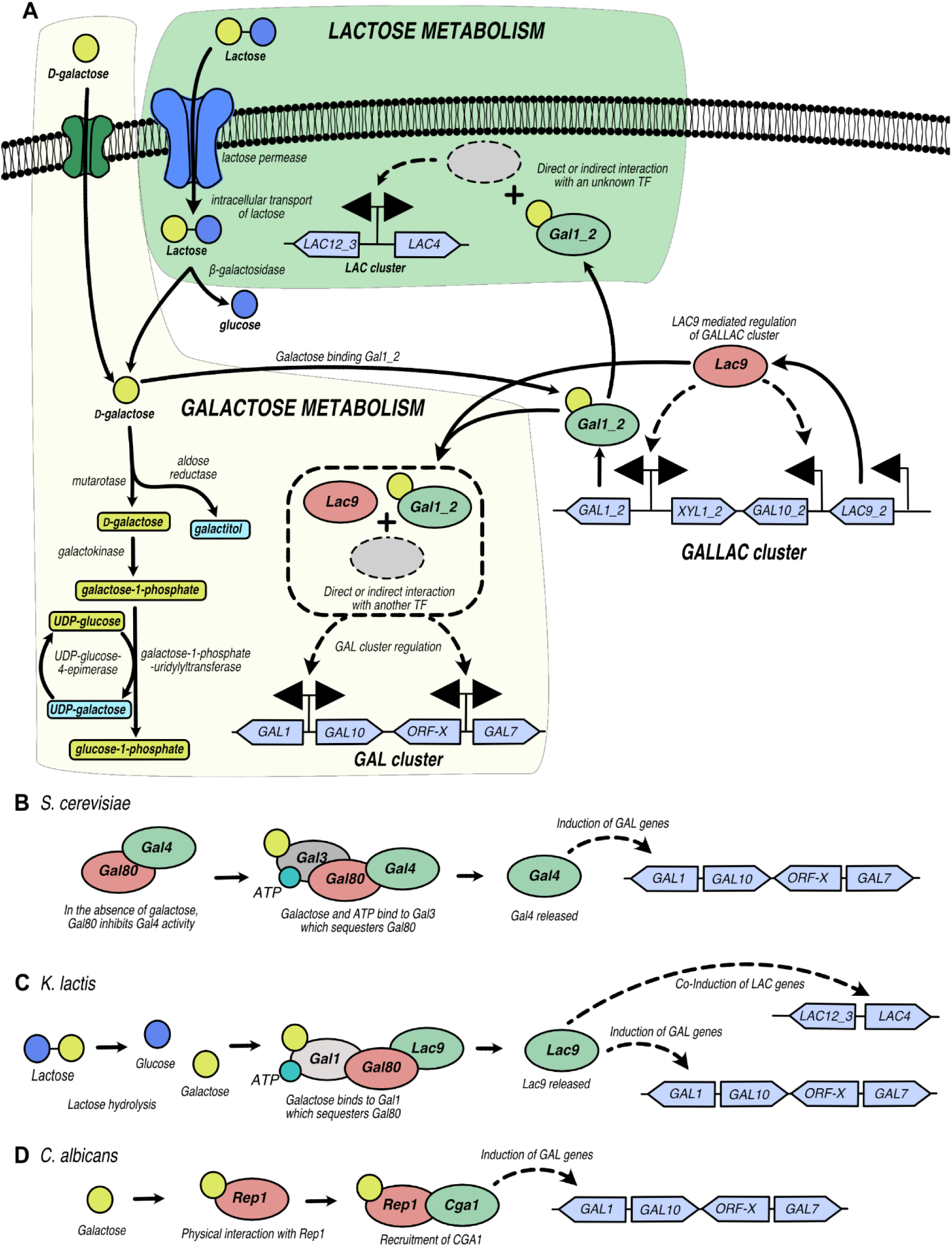
Graphic representation of regulatory mechanisms in *C. intermedia* and other yeast species: A) Depiction of lactose (green box) and galactose (light yellow box) metabolism in *C. intermedia* with the regulation of *GALLAC* cluster by the transcription factor CiLac9. On galactose, Lac9 and Gal1_2 interact directly or indirectly resulting in the regulation of *GAL* cluster gene(s), thus, affecting *C. intermedia’s* growth. On lactose, our results show that Gal1_2 from the *GALLAC* cluster regulates the *LAC* cluster at a transcriptional level. This effect of Gal1_2 can be speculated to be indirect due to the inability of Gal1_2 to bind DNA or protein based on predicted structure. Graphical representation also illustrates the overflow metabolism in *C. intermedia* because of aldose reductase mediated conversion of galactose to galactitol. B) Regulation of galactose metabolism in *S. cerevisiae* by the Gal3-Gal80-Gal4 system where galactose and ATP induce Gal3 to bind Gal80 resulting in the activation of Gal4. Thus, Gal4 induces structural GAL genes. C) Regulation of (ga)lactose genes in *K. lactis* is mediated by the bi-functional KlGal1. The ScGal4 homolog in *K. lactis* (KlGal1) is induced by galactose (or galactose derived from lactose) resulting in sequestering Gal80 and relieving Gal4 homolog, Lac9, which in turn activates the interconnected galactose and lactose metabolic genes in this yeast. C) Graphic representation of the Rep1 and Cga1 mediated galactose regulatory system in *C. albicans.* Galactose physically binds to Rep1 resulting in recruitment of Cga1 and the complex ultimately induces the structural genes responsible for galactose metabolism in this yeast.

Instead, we examined the role of Gal1 and Gal1_2 as regulators of lactose metabolism by performing β-galactosidase assays with *C. intermedia gal1Δ*, *gal1_2Δ* as well as WT (positive control) and *lacΔ* (negative control). Our RNAseq data showed that in WT, *LAC4* is expressed during growth on both galactose and lactose, respectively (Figure 2). Thus, we assessed the lactase activity during growth on both these sugars to include at least one condition where all strains could grow. For both galactose and lactose, lactase activity was readily detected in WT and *gal1Δ* cells but close to zero in the *lacΔ* and *gal1_2Δ* mutants (Figure 6C), showing that Gal1_2 is essential to induce lactase activity. Moreover, qPCR analysis of WT and *gal1_2Δ* showed that *LAC4* expression was diminished in *gal1_2Δ* as compared to the WT, indicating that regulation is exerted on the transcriptional level. In the same mutant we also observed that *GAL1* was still expressed (Figure 6D), fortifying the growth phenotyping results where we saw a clear difference in growth on galactose (+) and lactose (-) for this single mutant. Overall, these results firmly establish a difference in function between Gal1 and Gal1_2, where lack of Gal1_2 diminishes lactase transcription and activity while Gal1 does not, and further indicate important differences in regulation of lactose and galactose metabolism and growth.

## Discussion

In this work we have investigated how (ga)lactose is metabolized in the non-conventional yeast *C. intermedia* and shed light on the genetic determinants behind this trait. Interestingly, we found that the genome of *C. intermedia* contains not only the conserved *GAL* and *LAC* clusters, but also a unique *GALLAC* cluster that has evolved through gene duplication and divergence. By combining results from comparative genomics, transcriptomics analysis, deletion mutant phenotyping and metabolite profiling, we have started to unravel parts of the regulatory networks and interdependence of the three clusters and can show that the *GALLAC* cluster plays a vital role in both galactose and lactose metabolism in this yeast. With the Leloir pathway of budding yeasts acting like a model system for understanding the function, evolution and regulation of eukaryotic metabolic pathways, this work adds interesting new pieces to the puzzle.

Our results show that *C. intermedia* grows relatively fast on lactose, and strains of this species have been isolated several times from lactose-rich niches including fermentation products like white-brined cheese^46^ and cheese whey^47^. In these lactose-rich environments, survival likely necessitates a genetic makeup that can help outcompete rivaling microorganisms. Is the *GALLAC* cluster facilitating the fast lactose growth observed for *C. intermedia*, and if so, how? This is currently unresolved, but the genes within the cluster and the mutant phenotyping results provide some clues. First, the *GALLAC* cluster seems to have important regulatory functions, which can help to finetune metabolic fluxes and growth. We demonstrate that the cluster-encoded transcription factor Lac9_2 is important for onset of (ga)lactose growth, as deletion of *LAC9_2* leads to increased lag phase on both carbon sources. However, as *lac9_2Δ* cells eventually grow, Lac9_2 cannot be solely responsible for expression of the metabolic genes. Moreover, Lac9 binding motifs were only found in the promoters of *GALLAC* genes, suggesting that other transcriptional activators are responsible for induction of the *GAL* and *LAC* cluster genes.

In addition to Lac9_2, Gal1_2 from the *GALLAC* cluster also seems to be an important regulator of (ga)lactose growth. The bioinformatic analysis strongly suggests that *GAL1_2* in *C. intermedia* formed through gene duplication and divergence from the *GAL1* gene in the *GAL* cluster. Our results also show that Gal1_2 is essential for *LAC4* transcription and in extension, lactase activity and lactose growth, whereas deletion of *GAL1_2* alone did not abolish *GAL1* expression and galactose growth. Combined, these results indicate that the original Gal1 seems to have maintained the function as main galactokinase while Gal1_2 has taken on the role as a regulator. This evolutionary trajectory mirrors the path taken by Gal1 and Gal3 in *S. cerevisiae*^45^, but with a crucial distinction: the Gal1 proteins in *C. intermedia* have evolved in response to both lactose and galactose. On galactose, an additional deletion of *LAC9_2* was needed to impair growth, suggesting that the yeast senses and regulates expression of the galactose and lactose genes somewhat differently. Since Gal1_2 does not have a DNA binding capacity, we hypothesize that Gal1_2 binds galactose and thereafter activates unknown transcription factor(s) that ultimately bind and induce expression from the *LAC* and *GAL* clusters. Although many details are still to be elucidated, it is clear that *C. intermedia* has developed a way of regulating its (ga)lactose metabolism that differs from other yeast species studied to date, including the Gal3-Gal80-Gal4 regulon in *S. cerevisiae*^48^, the Gal1-Gal80-Lac9 equivalent in *K. lactis*^40^ and the Rep1-Cga1regulatory complex in *C. albicans*^29^. Future research will include identifying these unknown TFs and fully elucidating the roles of Lac9_2 and Gal1_2 in sensing, signaling, and regulating the cellular response to changes in the nutritional environment.

Another interesting feature of the *GALLAC* cluster is the *XYL1_2* gene encoding an aldose reductase. Although no galactitol or other intermediates of an oxidoreductive pathway accumulate in the WT under the growth conditions assessed, several of the constructed mutants (in particular, *galΔ* and *gal1Δ*) accumulate galactitol upon growth on lactose. In *S. cerevisiae,* galactitol functions as an overflow metabolite ensuring that cells avoid accumulation of galactose-1-phosphate, a known toxic intermediate of the Leloir pathway in the cell^15,20^, and it is reasonable to assume that the same is true for *C. intermedia.* Moreover, it is interesting to note that aldose reductases can directly convert *β*-D-galactose, the hydrolysis product of lactose, whereas galactokinase requires *β*-D-galactose conversion into *α*-D-galactose before it can be metabolized via the Leloir pathway. We speculate that induction of an aldose reductase gene in tandem with the *LAC* and *GAL* genes in response to lactose (and galactose) can be an efficient way to quickly metabolize these sugars, providing a growth advantage in competitive lactose-rich environments.

In addition to the basic scientific questions that can be answered by studying evolution and sugar metabolism in lactose-growing yeast species, these yeasts can also be used as cell factories in industrial biotechnology processes. Here, a better understanding of the underlying genetics for this trait enables metabolic engineering to optimize the conversion of lactose-rich whey into value-added products. The dairy yeasts *K. lactis* and *K. marxianus* have been developed and used for whey-based production of ethanol^49^, recombinant proteins^50^ as well as bulk chemicals such as ethyl acetate^51^, while exploration of new lactose-metabolizing yeasts allows for additional product diversification. With lactose as substrate, a carbon-partition strategy can be used for bioproduction, where the glucose moiety is converted into energy and yeast biomass and the galactose moity in steered into production of the wanted metabolite, or vice versa^13^. Through this strategy, the non-conventional yeast *C. intermedia* can also be explored to produce various growth-coupled metabolites, including galactitol and derivatives thereof.

In conclusion, our work on the non-conventional, lactose-metabolizing yeast *C. intermedia* has paved the way towards a better understanding of the (ga)lactose metabolism in this relatively under-studied species. To the best of our knowledge, we show for the first time that gene duplication and divergence resulted in the formation of a unique *GALLAC* cluster and its essential role in (ga)lactose metabolism in this yeast, providing new insights of how organisms can evolve metabolic pathways and regulatory networks. In addition, the proven ability of *C. intermedia* to grow relatively well on lactose establishes this yeast as an interesting lactose-assimilating species also for future industrial applications.

## Materials and Methods

### Culture conditions and molecular techniques

For amplification of plasmids, *E. coli* was grown on LB medium (1 % tryptone, 1 % NaCl and 0.5 % yeast extract) containing ampicillin (100 μg/mL) for plasmid selection.

*C. intermedia* CBS 141442 was grown in YPD medium (1% yeast extract, 2% bactopeptone and 2% glucose) prior to yeast transformation using the split marker technique as described previously^39^. Using this technique, deletion cassettes were constructed as two partially overlapping fragments, each containing half of the selection marker fused to either upstream or downstream sequences of the target gene. Deletion fragments were transformed using electroporation (BioRad Micropulse electroporator). After transformation, cells were plated on YPD agar containing 200 μg/ml nourseothricin to select for integration and expression of the *CaNAT1* selection marker.

Colony PCR was used to identify transformants with correct gene deletions, where single colonies were resuspended in 50 μL dH_2_O using a sterile toothpick and then heated to 90 °C for 10 min. After cooling to 12 °C, 2 μL of each suspension was used as a template for PCR using PHIRE II polymerase (Thermo-Fisher Scientific, USA). For each mutant, three PRC primers were used, where the first primer was designed to hybridize to the genome outside the flanking region, the second to the marker gene and the third to the targeted gene (negative control). For each gene deletion, three correctly targeted transformants were selected for subsequent phenotyping.

To construct the double gene deletion mutant (lac9_2, the split marker method was used twice in the same strain background, first employing the split *CaNAT1* selection marker as described above, and then a split KanMX selection marker PCR amplified from the plasmid pTO149_RFP_CauNEO developed for *Candida auris*^52^. Correctly assembled and genome integrated KanMX markers gave rise to *C. intermedia* transformants resistant to the antibiotic Geneticin (200 μg/mL).

For complementation tests in *S. cerevisiae*, *C. intermedia GAL1* and *GAL1_2* genes were synthesized and cloned in a vector backbone (pESC-URA; GenScript Biotech, New Jersey, USA). Codon CTG were adjusted to alternate codon prior to optimization of the complete gene for expression in *S. cerevisiae* using the GenSmart™ Codon Optimization tool (GenScript Biotech, New Jersey, USA). *S. cerevisiae* BY4741/2 *GAL1* knockouts used for complementation experiment were grown on YP media with 2% glucose and transformed with above mentioned plasmids using LiAc/PEG heat-shock method^53^. Transformants were selected on agar plates with YNB-uracil and 2% glucose, restreaked and then tested for growth in liquid YNB-URA media with 2% galactose in GrowthProfiler at 30 °C and 250 rpm. *S. cerevisiae* BY4741/2 *gal1Δ* transformed with p426 (empty vector with *URA3* as selection marker) was used as negative control.

### Growth Experiments

#### Growth Profiler

To follow growth over time for *C. intermedia* CBS 141442 and the other yeasts characterized in this work, strains were precultured at 30 °C, 180 rpm overnight in synthetic defined minimal Verduyn media^54^ containing 2% glucose (w/v). Precultured cells were then inoculated in 250 µL minimal media supplemented with 20 g/L carbon source to a starting OD_600_ = 0.1. All yeast strains were grown in biological triplicates in a 96-well plate setup in a GrowthProfiler 960 (Enzyscreen, Netherlands). ‘Green Values’ (GV) measured by the GrowthProfiler correspond to growth based on pixel counts, and GV changes were recorded every 30 min for 72 h at 30 °C and 150 rpm.

#### Cell growth quantifier (CGQ)

Growth characterization was also performed in shake flasks using Cell Growth Quantifier (CGQ-Scientific Bioprocessing, Germany) ^55^. Wild type and mutant strains were precultured at 30 °C, 200 rpm overnight in synthetic defined minimal Verduyn media containing 2% glucose (w/v), followed by inoculation of 25ml of minimal medium supplemented with 2% carbon source (galactose or lactose) in 100mL shake flasks to a starting OD_600_ = 0.1. Growth was quantified as “Scatter values” by the CGQ system^56^. Scatter values were recorded for 10 days at 30 °C and 200 rpm for each strain growth in biological triplicates and sampling was performed for sugar and polyol analysis.

### Lactase activity assay

β-galactosidase activity was determined using the Yeast β-Galactosidase Assay Kit (Thermo-Fisher Scientific, USA) following the manufacturer’s instructions. Cells were harvested at different timepoints during growth and tested for lactase activity. A Working solution was prepared by mixing equal amounts of 2X β-galactosidase Assay Buffer (containing ortho-nitrophenyl-β-galactoside (ONPG)) and Yeast Protein Extraction Reagent. The reaction was initiated by mixing 100uL of working solution with 100uL cell culture and incubated for 30 min at 37 °C in a thermomixer. After 30 min, cell mix was centrifuged at 5000 rpm for 3 mins and the supernatant was analyzed for lactase activity by measuring o-nitrophenol release from ONPG at 420 nm in microplate reader (FLU-Ostar Omega-BMG LabTech, Ortenberg, Germany).

### Determination of sugar and polyol concentrations

Sugars and galactitol concentrations were measured using a Dionex high-performance liquid chromatography (HPLC) system equipped with an RID-10A refractive index detector and an Aminex HPX-87H carbohydrate analysis column (Bio-Rad Laboratories). Analysis was performed with the column at 80 °C, and 5 mM H_2_SO_4_ as mobile phase at a constant flow rate of 0.8 mL/min. Culture samples were pelleted prior to analysis, following which, the supernatant was passed through a 0.22 μm polyether sulfone syringe filter. Chromatogram peaks were identified and integrated using the Chromeleon v6.8 (Dionex) software and quantified against prepared analytical standards.

### Comparative genomics and evolutionary mapping

We established the blast database for 332 yeast species based on the work of Shen et al., 2018^1^. Then we used tblastn to get gene hits for each specific gene in three clusters against 332 yeast species. Based on the generated data, we further mapped gene hits from species to clade levels. To investigate the evolution of genes in the *GALLAC* cluster, a comprehensive pipeline based on the work of Goncalves and colleagues was developed^57^. For each candidate gene in the *GALLAC* cluster, BLASTP was run against the NCBI non-redundant (nr) protein sequence database and homologs were selected according to the top 300 BLAST hits to each query sequence. These homologs were aligned with `MAFFT v7.310`^58^ using default settings for multiple sequence alignment. Poorly aligned regions were removed with `trimAl`^59^ using the ‘-automated1’ option. Subsequently, phylogenetic trees were built using `IQ-TREE v1.6.12` ^60^ with 1000 ultrafast bootstrapping replicates^61^. Each tree was rooted at the midpoint using a customized script combining R packages `ape v5.4-1` and `phangorn v2.5.5.` Finally, the resulting phylogenies were visualized using `iTol v5`^62^.

### Transcription factor binding motif analysis

To determine the binding motifs of transcription factors in promoter regions of the *GAL*, *LAC* and *GALLAC* cluster, MEME (Version 5.5.43) promoter binding motif analysis was used. Promoter regions of all genes from the three clusters were added as query sequences with the following constraints: maximum number of motifs = 5, maximum length of motif = 25 bases, any number of motif repetitions (-anr), background model = 0-order model of sequences. Motif(s) derived from this analysis were then fed as input to Tomtom^63^ (version 5.5.4) to compare against Yeastract^64^ database.

### RNA sequencing

Transcriptomics using RNA sequencing was performed as previously described^37^. In brief, *C. intermedia* CBS 141442 was grown in controlled stirred 1-L bioreactor vessels (DASGIP, Eppendorf, Hamburg, Germany) containing 500 mL synthetic defined minimal Verduyn media with 2% Glucose, Galactose or Lactose. Reactor conditions were maintained as: Temp = 30 °C; pH = 5.5 (maintained with 2M Potassium Hydroxide); Aeration = 1 Vessel Volume per Minute; stirring = 300 rpm.

#### RNA extraction

For RNA extraction, samples (10 mL) were collected when the dissolved oxygen of the culture was 35–40% (v/v). After washing the cells, the pellets were immediately frozen using liquid nitrogen. Frozen pellets were stored at −80 °C until extraction. The frozen pellets were thawed in 500 μL of TRIzol (Ambion—Foster City, CA, USA) and thoroughly resuspended. Then, cells were lysed in 2 mL tubes with Lysing Matrix C (MP Biomedical, Santa Ana, CA, USA) in a FastPrep FP120 (Savant, Carlsbad, CA, USA) for five cycles, at intensity 5.5 for 30 s. Tubes were cooled on ice for a minute between cycles and resuspended once again in 500 μL of TRIzol and vortexed thoroughly. After incubation at room temperature for 5 min, tubes were centrifuged for 10 min at 12,000 rpm and 4 °C. Chloroform was added to the collected supernatants (200 μL of chloroform per mL of supernatant) and vortexed vigorously for 30 s. After centrifugation for 15 min at 12,000 rpm, 4 °C, the top clear aqueous phase was collected and transferred to a new RNase-free tube, to which, equal amount of absolute ethanol was slowly added while mixing. Each sample was loaded into a RNeasy column (RNeasy Mini Kit, Qiagen—Hilden, Germany) and further steps followed the protocol of the manufacturer. The RNA was eluted with RNase-free water and samples were stored at −80 °C until use.

#### Data analysis

RNA samples were analyzed in a TapeStation (Agilent, Santa Clara, CA, USA), and only samples with RNA integrity number above 8 were used for library preparation. Sequencing using the HiSeq 2500 system (Illumina Inc.—San Diego, CA, USA), with paired-end 125 bp read length, and v4 sequencing chemistry, was followed by quality control of read data using the software FastQC version 0.11.5^65^. Software Star version 2.5.2b^66^ was used to map reads to the reference genome. Gene counts were normalized with weighted trimmed mean of M-values using the calcNormFactor function from the package edgeR^67^ and Limma package^68^ were used to transform and make data suitable for linear modelling. The estimated p-values were corrected for multiple testing with the Benjamini-Hochberg procedure, and genes were considered significant if the adjusted p-values were lower than 0.05. The raw counts were filtered such that genes with CPM > 3.84 in at least 12% (5/43) of the samples were retained. The R function ‘varianceStabilizingTransformation()’ from R package ‘DESeq2’^69^ was used to convert raw counts to variance-stabilized-counts (VST). Expression data for *C. intermedia* on galactose and lactose was normalized using glucose as control condition. The RNA seq datasets are available in the European Nucleotide Archive (ENA) with the accession number E-MTAB-6670.

### Gene expression analysis using qPCR

Primers used for mRNA quantification using qPCR are listed in Table S1. Primers were designed using Primer3 (https://primer3.ut.ee/) and were checked for efficiency. Only primers having efficiency between 90-110% were used for qPCR. Cultures were grown at 30 °C and 200 rpm in 100 ml shake flasks containing 25 ml synthetic defined minimal Verduyn media containing either 2% glucose (control), galactose or lactose as carbon source. Cells were harvested for each strain at lag, early log and late log phases, taking three biological replicates. Harvested cells were pelleted by centrifugation at 4 °C for 5 mins at 5000 rpm and washed twice by resuspending in ice-cold sterile dH_2_O water and centrifugation. Cell pellet was snap-frozen using liquid nitrogen and stored at −80 °C for cDNA synthesis. RNA extraction was performed as described for RNA sequencing above. cDNA synthesis and RT qPCR analysis was performed using Maxima H Minus First Strand cDNA Synthesis Kit (Thermo Fisher) and Maxima SYBR Green/Fluorescein qPCR Master Mix (2X) (Thermo Fisher), according to the manufacturer’s instruction. Fold change was calculated using the delta-delta Ct method (2^-^*^ΔΔ^*^Ct^) with expression values in glucose as control condition and Ci*ACT1* as the reference gene for normalization.

## Acknowledgments

Authors would like to thank Peter Dahl from Department of Chemistry & Molecular Biology, Gothenburg University for providing *S. cerevisiae* BY4741 and BY4742 knockout strains and Dr. Xiang Jiao from Department of Life Sciences, Chalmers University of Technology for providing the plasmid *p426*. Authors would also like to thank ARS culture collection (NRRL) for providing us with different lactose metabolizing strains upon request.

## Author contributions

Conceptualization: K.V.R.P. and C.G.; methodology: K.V.R.P., L.Y., K.P., F.F.O., and C.G.; investigation: K.V.R.P., L.Y., K.P., F.F.O., J.L. and C.G.; original manuscript draft preparation: K.V.R.P. and C.G.; and manuscript review and editing: all.

## Availability of data

The RNA-Seq datasets are available in the in the European Nucleotide Archive (ENA) with the accession number E-MTAB-6670.

## Supplementary figures and figure legends

**Figure S 1:**
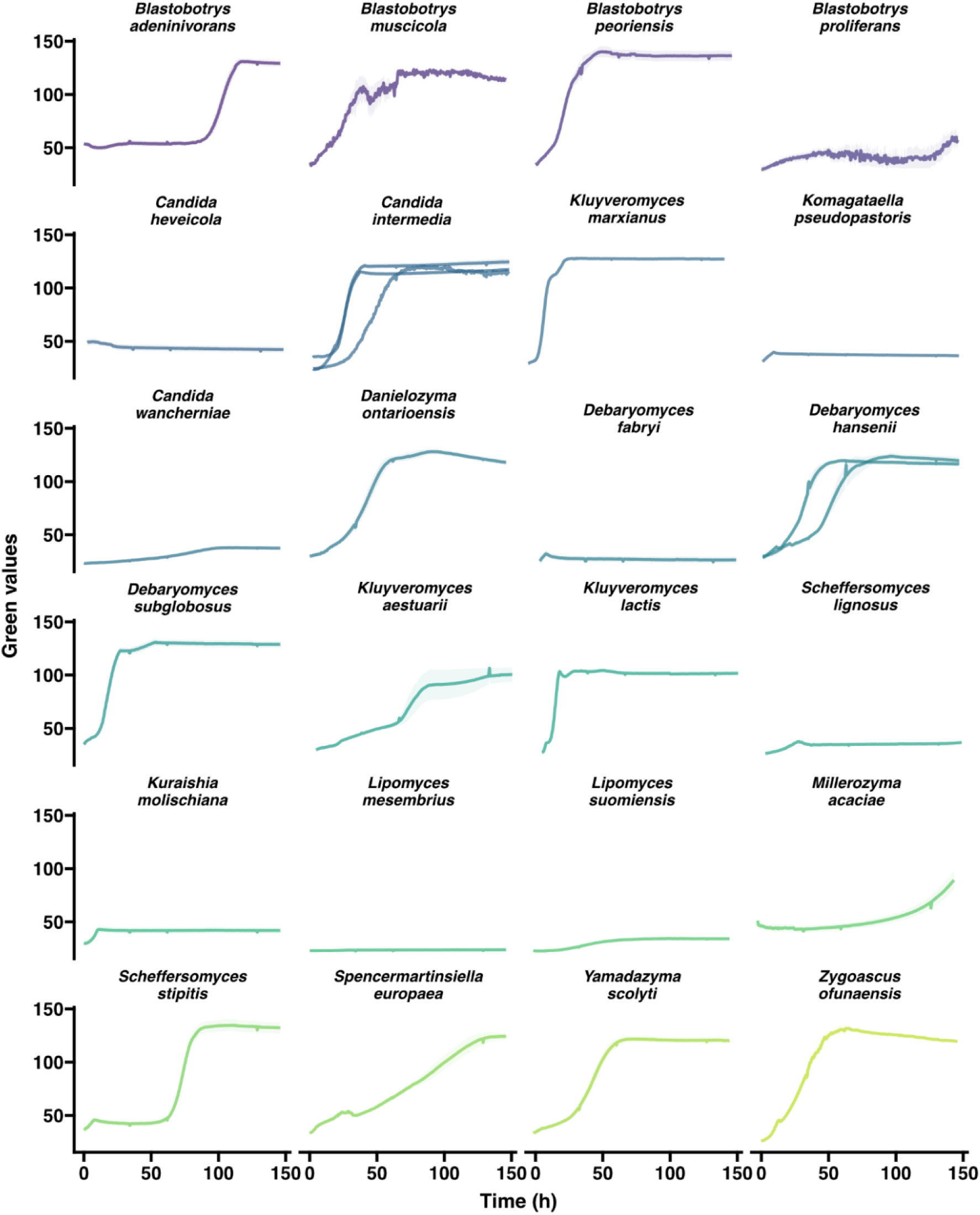
Growth curves for 24 lactose growing species from the work of Shen et al ^1^. The graphs depict data procured from GrowthProfiler in 96-well format, plotted as mean ± standard deviation (shaded region) for biological triplicates per strain. On y-axis final biomass yield is depicted in green values (G.V. - corresponding to growth based on pixel counts, as determined by a GrowthProfiler instrument) and is plotted against time (h) on x-axis.

**Figure S 2:**
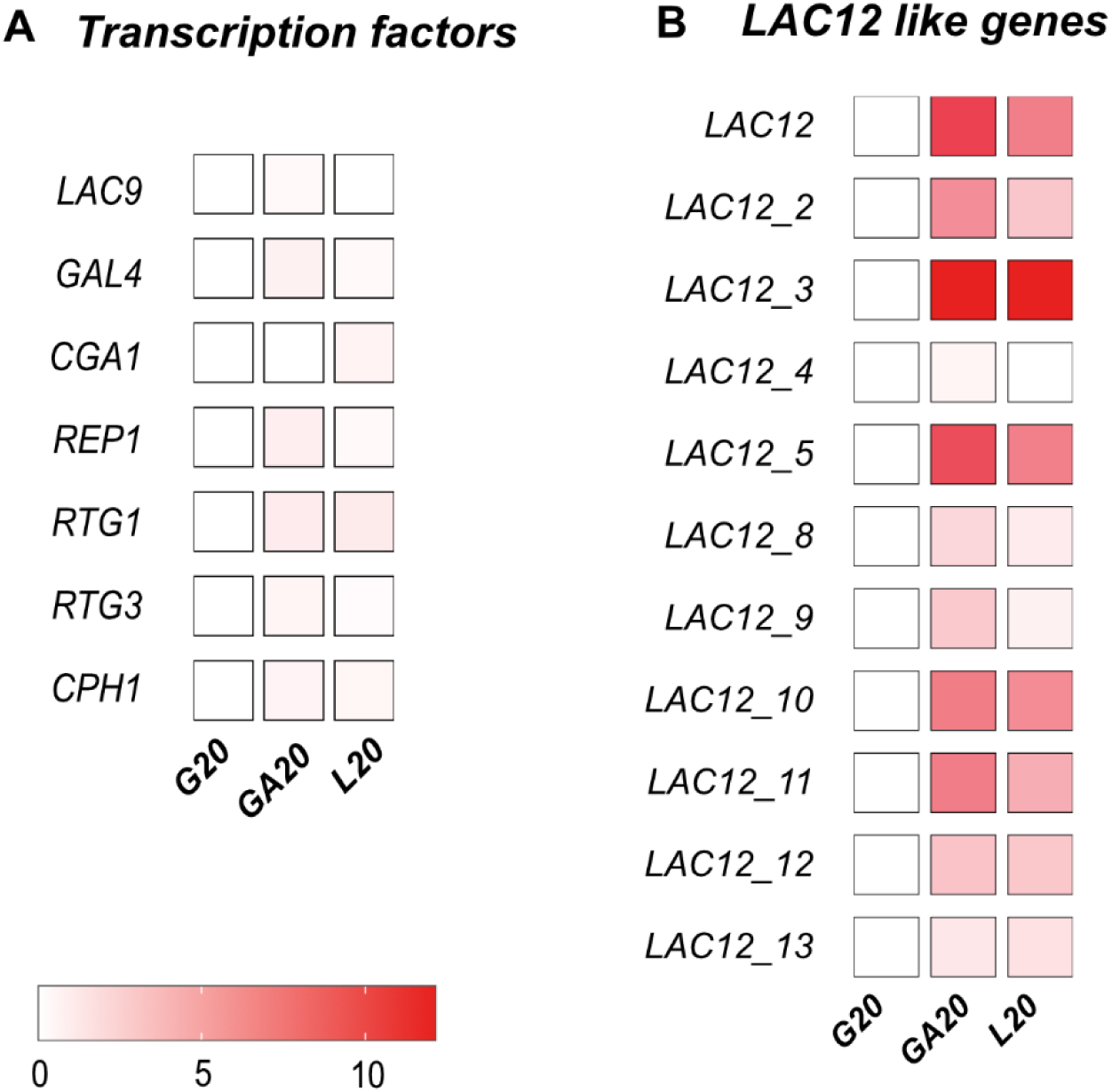
Gene expression pattern for different transcription factor orthologues and Lac12 like genes in *C. intermedia*. Gene expression in Galactose (GA20) and Lactose (L20) have been normalized for values on Glucose (G20). Legend shows expression (in fold change) from 0 to 10 in increasing gradient of red.

**Figure S 3:**
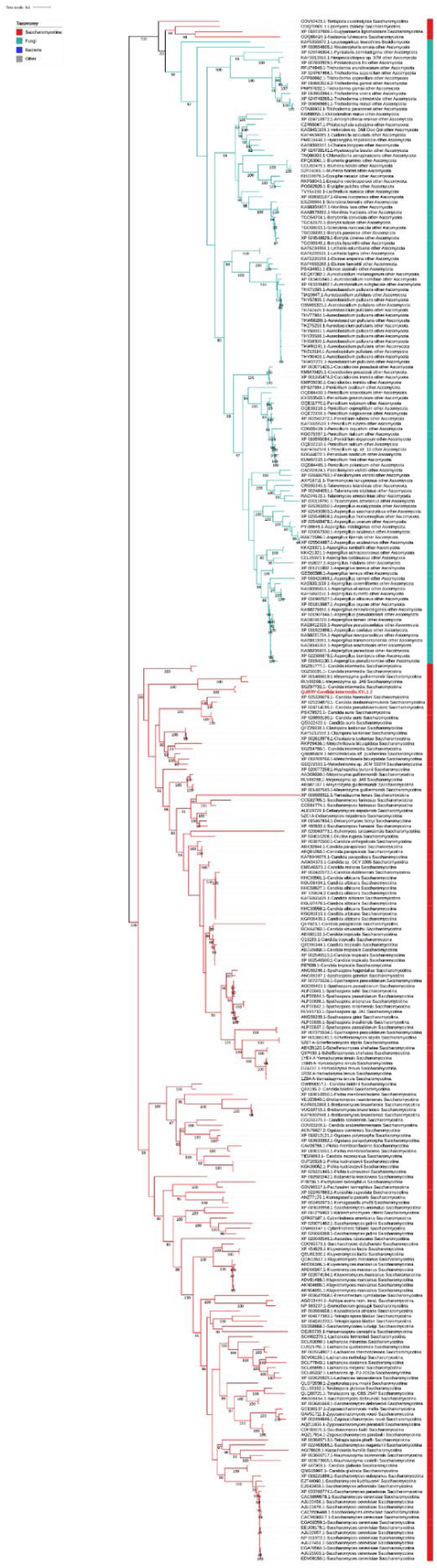
Maximum likelihood phylogenetic tree depicting the origin and evolution of the *XYL1_2* gene in *C. intermedia*.

**Figure S 4:**
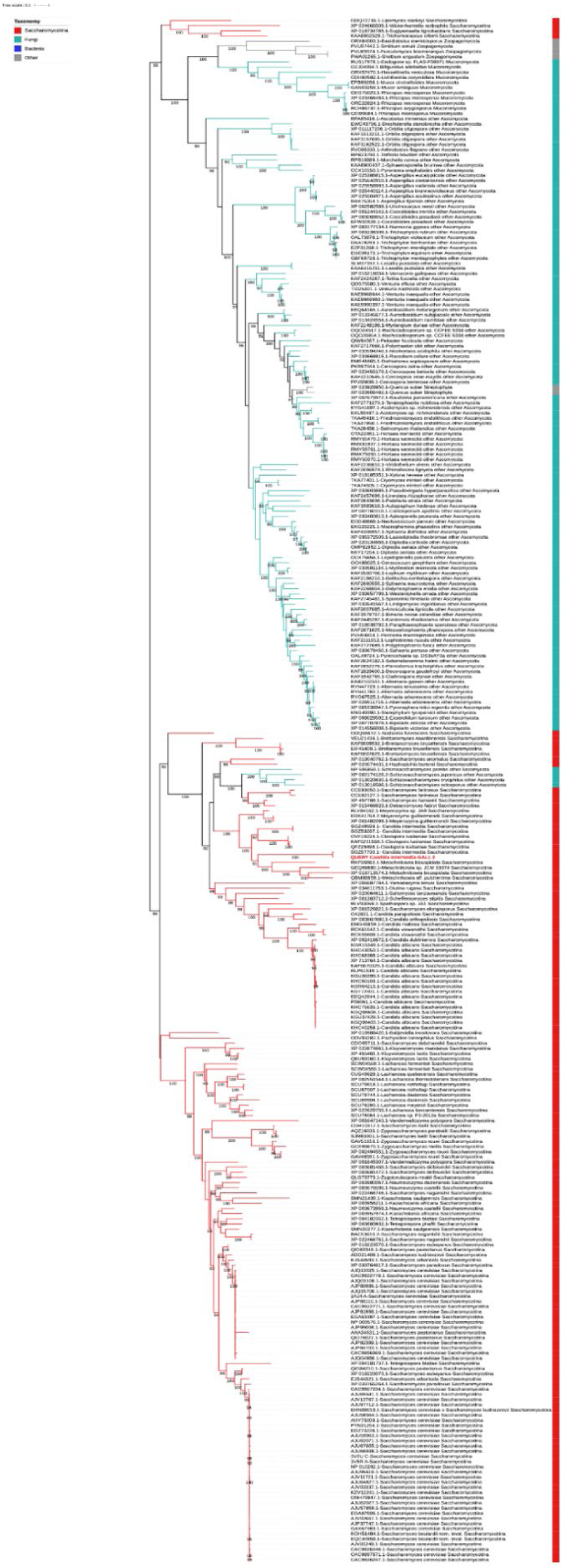
Maximum Likelihood phylogenetic tree depicting origin and evolution of the *GAL1_2* gene in *C. intermedia*.

**Figure S 5:**
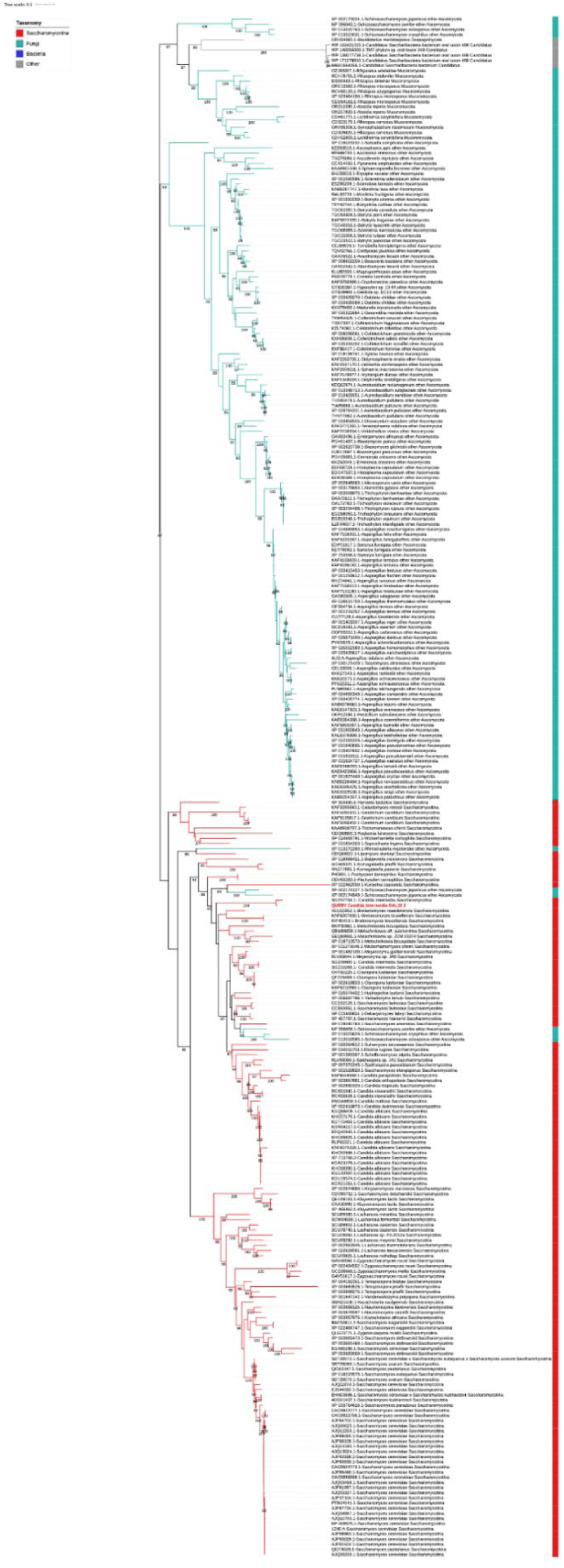
Maximum likelihood phylogenetic tree for the origin and evolution of the *GAL10_2* gene in *C. intermedia*.

**Figure S 6:**
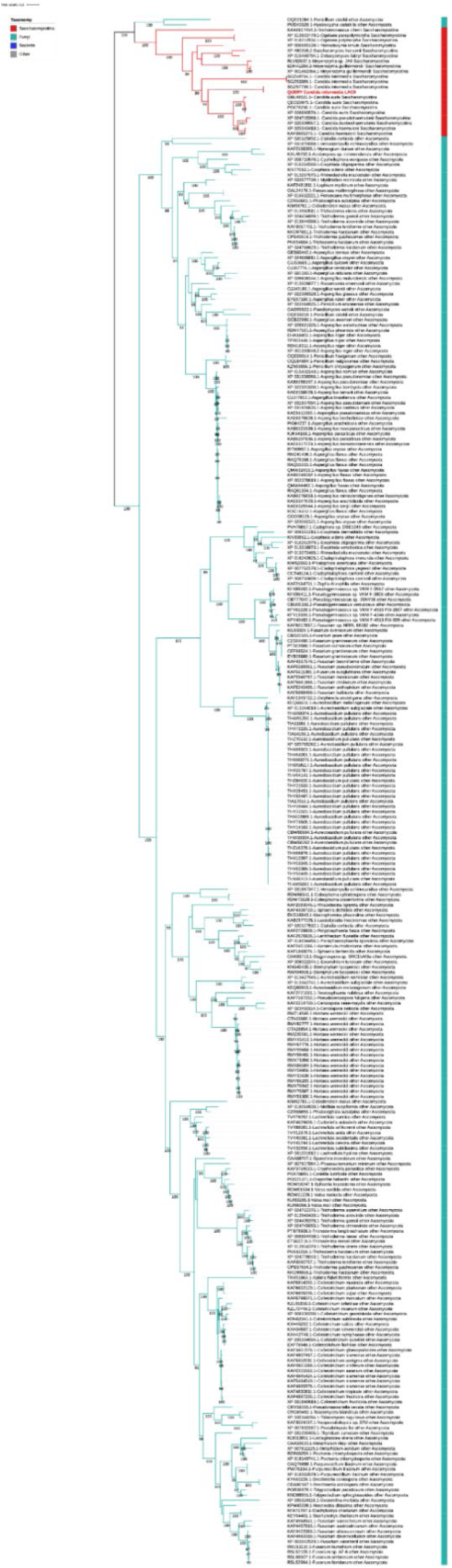
Maximum likelihood phylogenetic tree for the origin and evolution of the *LAC9* gene in *C. intermedia*.

**Figure S 7:**
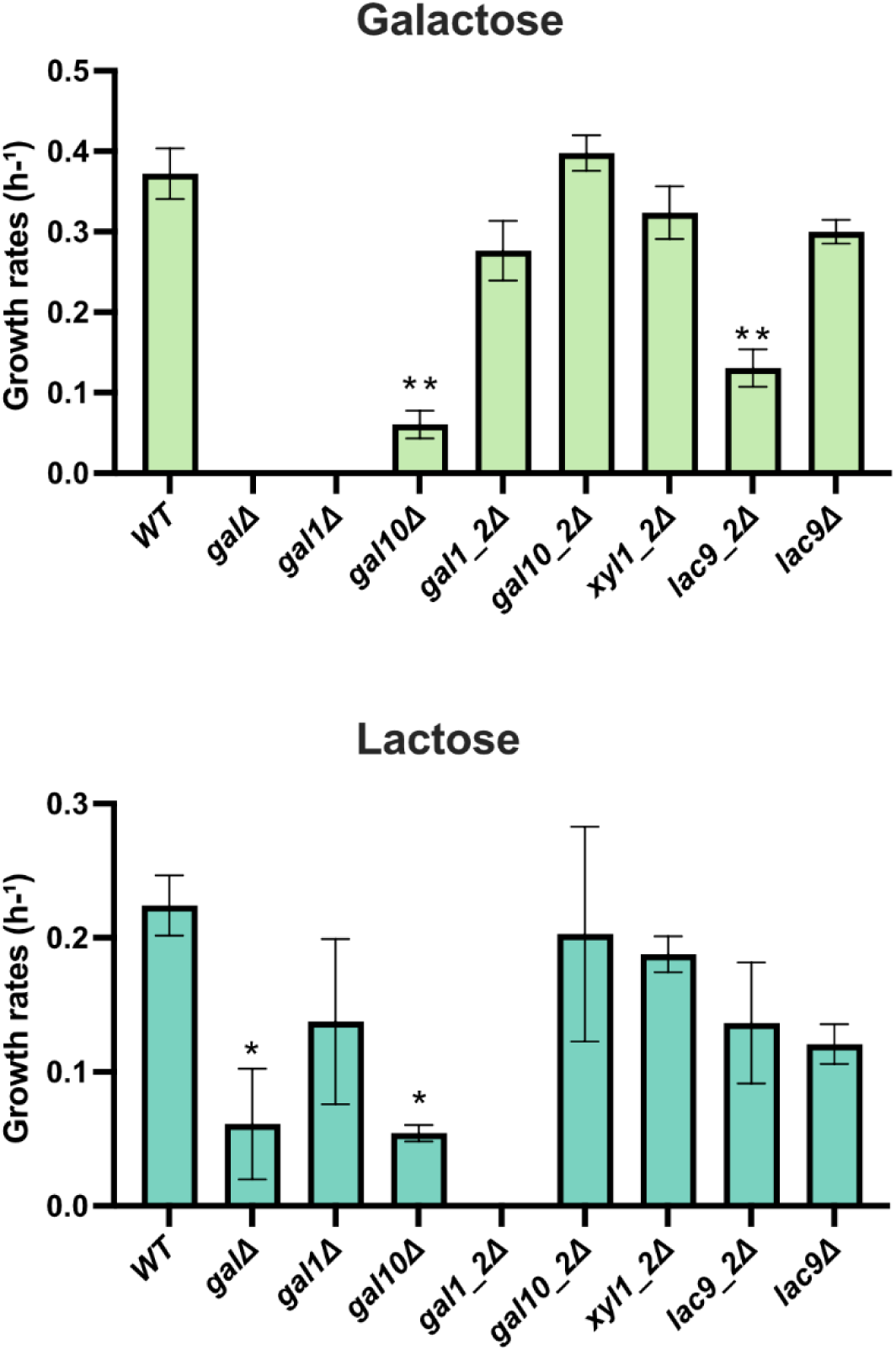
Bar plot with growth rates of different mutants in comparison to the wild-type strain (WT) on galactose as well as lactose. Significance difference in growth rates compared to the WT strain have been estimated using students t-test and values with p > 0.01 are considered significantly different. Data are represented as mean ± standard deviation for biological triplicates.

**Figure S 8:**
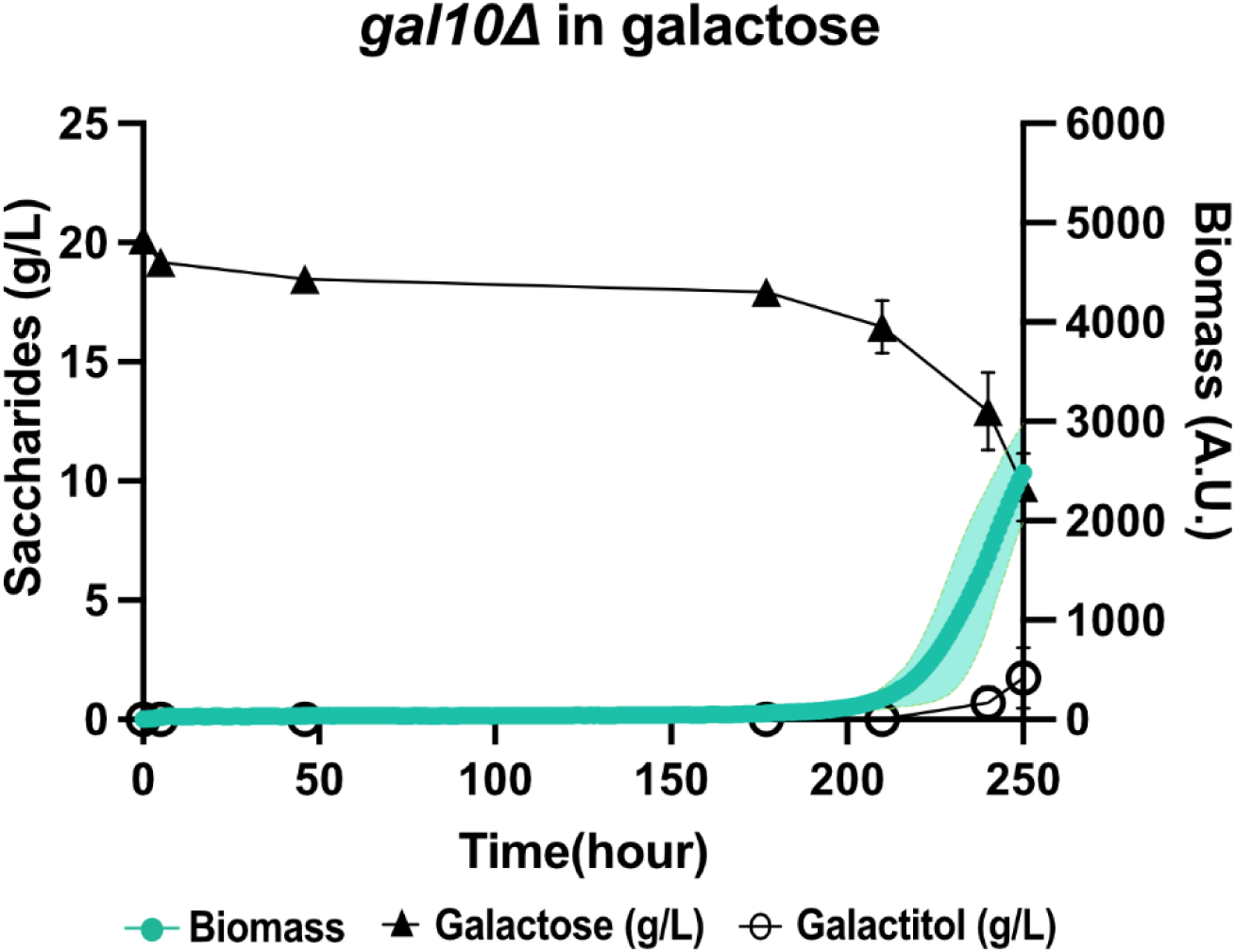
Growth and metabolite profile for gal10Δ in galactose containing minimal media. Graph represents biomass (filled green circle) on the right y-axis, consumption of respective sugars (filled triangle for galactose in g/L) and metabolite production (open circle for galactitol in g/L) on the left y-axis, plotted against time (in hours) on x axis. Data are represented as mean ± standard deviation for biological triplicates.

**Figure S 9:**
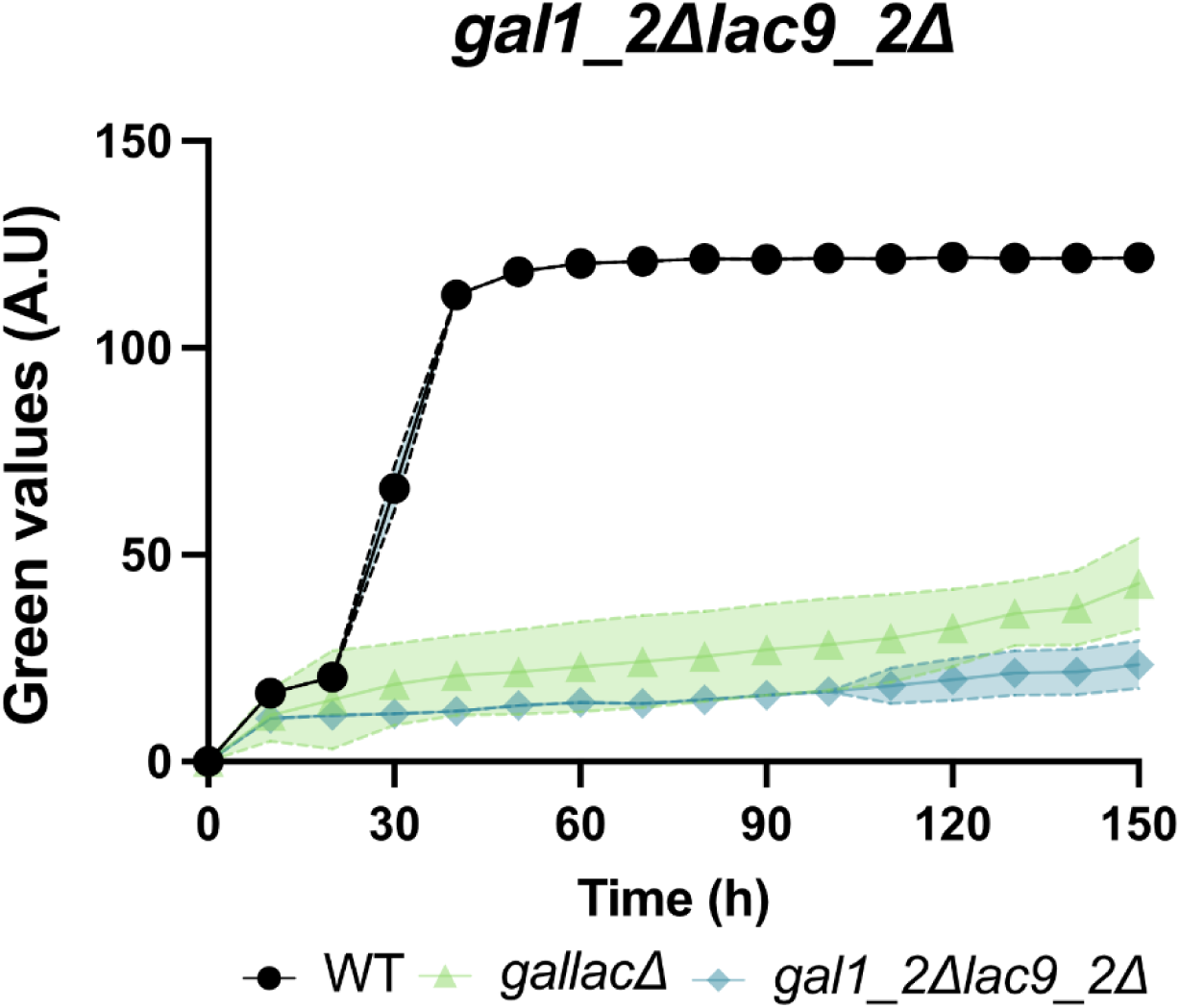
Growth profile for the double deletion mutant *gal1_2Δlac9Δ* in comparison with WT and *gallacΔ* in galactose. Legend shows the wild-type strain (black circle), *GALLAC* cluster deletion mutant (light green triangle), *gal1_2Δlac9Δ* depicted in the graph with growth as green values (A.U.) on the y-axis against time(hours) on the x-axis. Data are represented as mean ± standard deviation for biological triplicates indicated by colors: wild type – yellow, lac cluster mutant – purple, gallac cluster deletion – dark green and gal cluster mutant – light green.

**Figure S 10:**
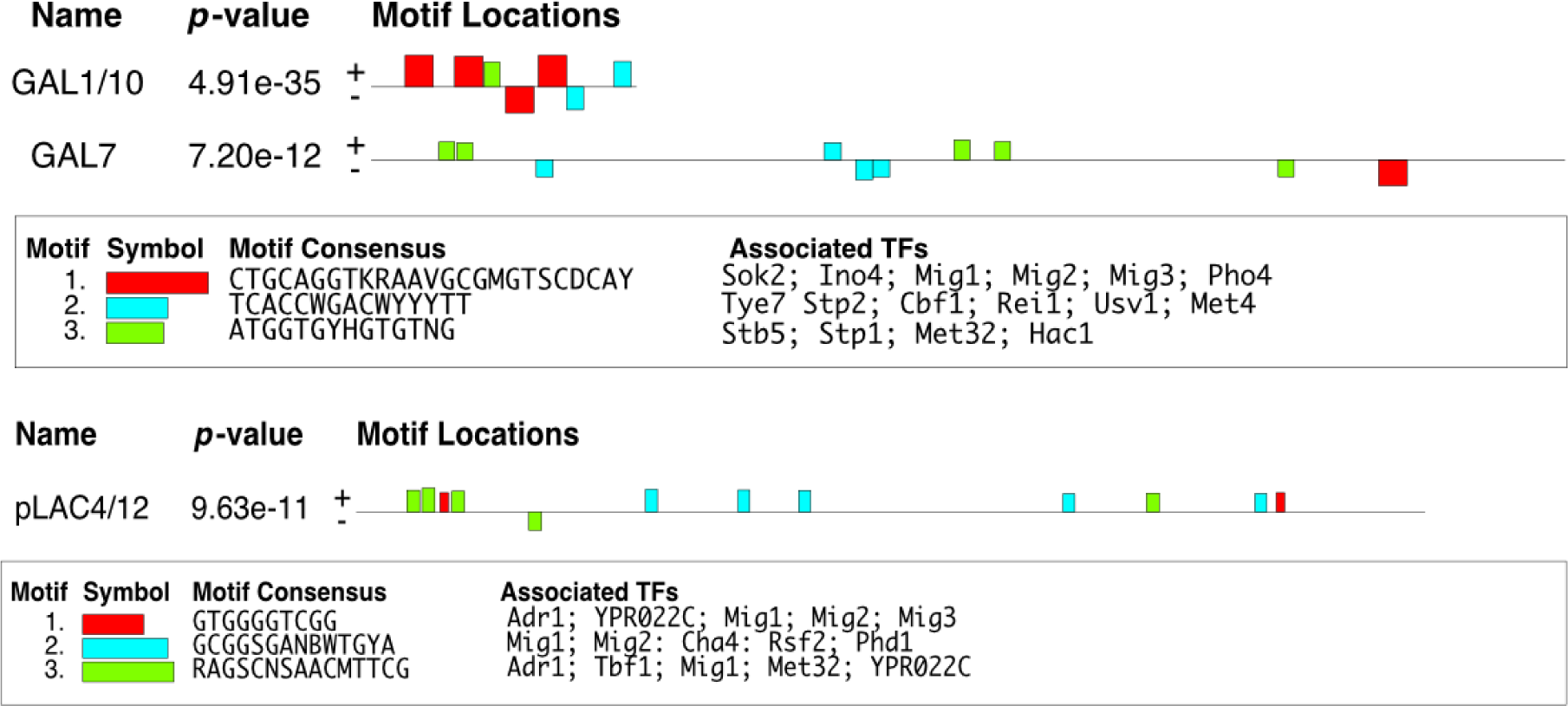
Results of transcription factor binding analysis performed using MEME (version 5.5.2) on promoter regions of genes in the *GAL* and *LAC* clusters. Results show the predicted binding motifs of TFs in the promoter regions ranked based on p-value for the motif. Also mentioned are the predicted transcription factors that are associated to the binding motifs.

**Figure S 11:**
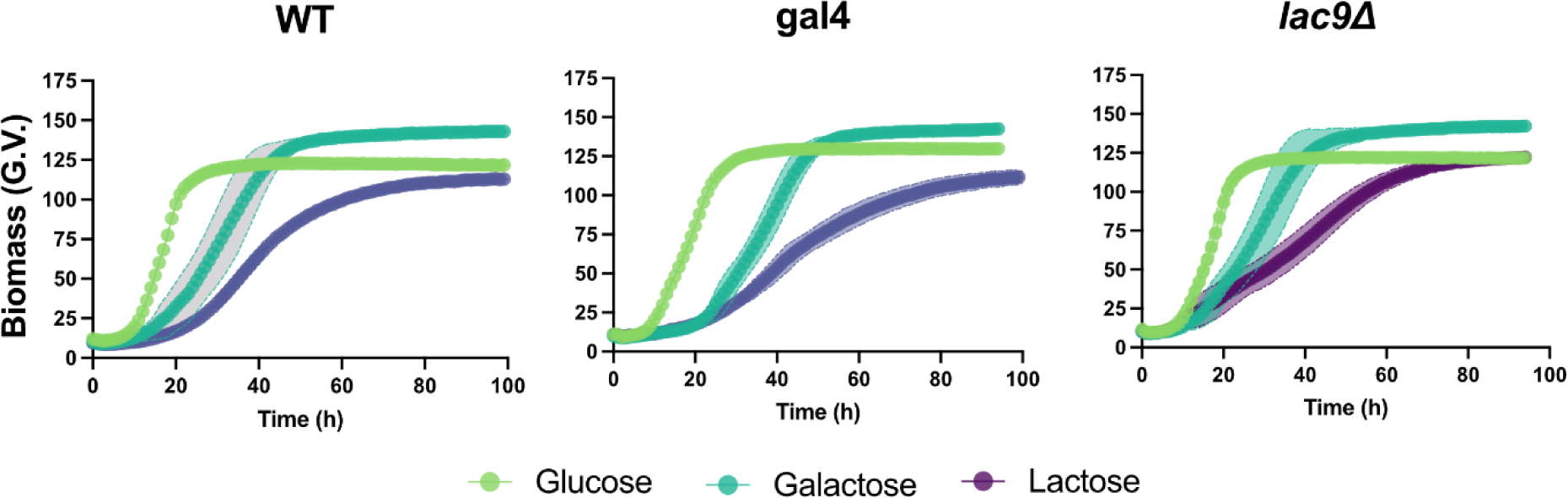
Growth profiles for WT, gal4 and lac9 mutants in glucose (light green), galactose (dark green) and lactose (purple) containing media. Time (in hours) on x-axis is plotted against biomass yield (green values – G.V.) on y-axis. Data are represented as mean ± standard deviation for biological triplicates.

**Table S 1:**
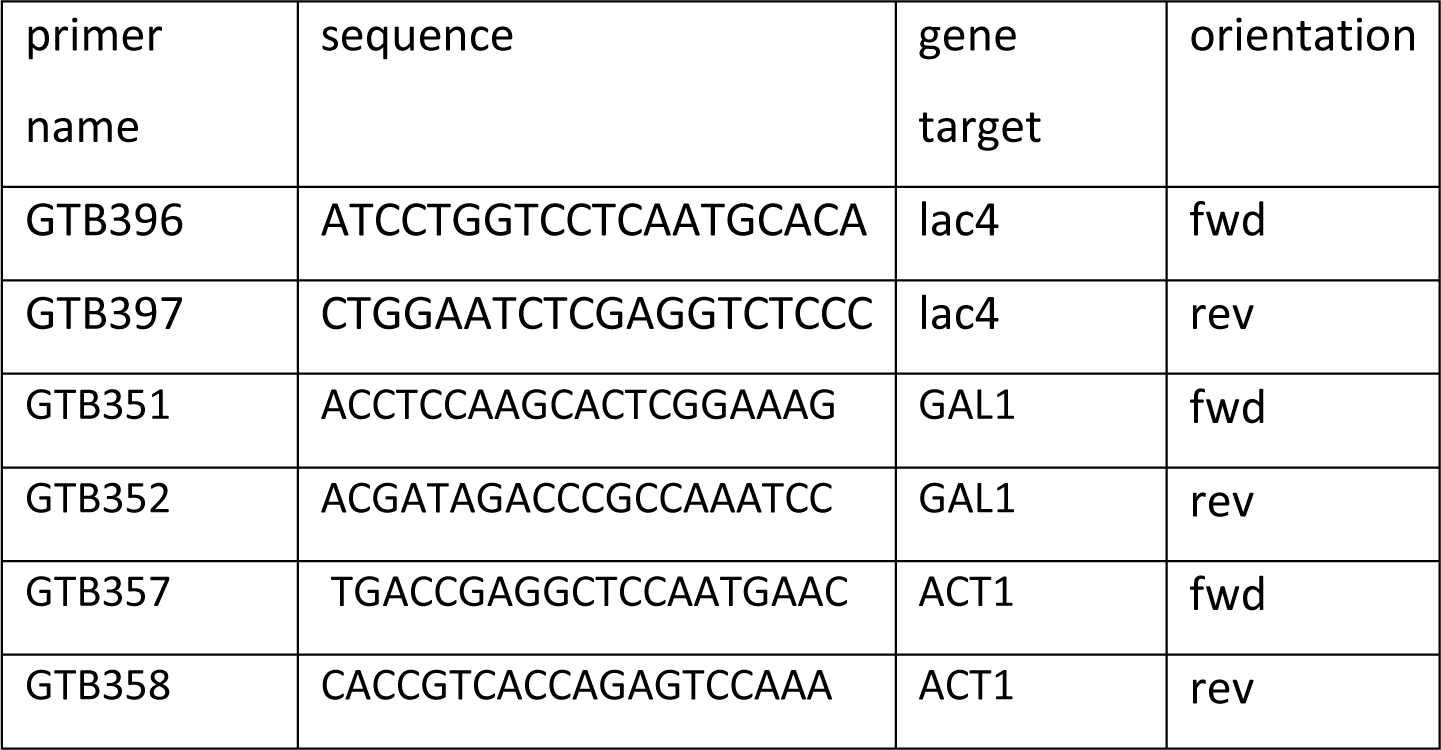
Primers used for mRNA quantification using qPCR in *C. intermedia*. Primers were designed using Primer3 (https://primer3.ut.ee/) and primer pairs were checked for efficiency prior to use.

## References

1. Shen, X.X., Opulente, D.A., Kominek, J., Zhou, X., Steenwyk, J.L., Buh, K.V., Haase, M.A.B., Wisecaver, J.H., Wang, M., Doering, D.T., et al. (2018). Tempo and Mode of Genome Evolution in the Budding Yeast Subphylum. Cell 175, 1533–1545 e1520. 10.1016/j.cell.2018.10.023.

2. Capuco, A.V., and Akers, R.M. (2009). The origin and evolution of lactation. J Biol 8, 37. 10.1186/jbiol139.

3. Schaffrath, R., and Breunig, K.D. (2000). Genetics and molecular physiology of the yeast Kluyveromyces lactis. Fungal Genet Biol 30, 173–190. 10.1006/fgbi.2000.1221.

4. Godecke, A., Zachariae, W., Arvanitidis, A., and Breunig, K.D. (1991). Coregulation of the Kluyveromyces lactis lactose permease and beta-galactosidase genes is achieved by interaction of multiple LAC9 binding sites in a 2.6 kbp divergent promoter. Nucleic Acids Res 19, 5351–5358.

5. Lane, M.M., Burke, N., Karreman, R., Wolfe, K.H., O’Byrne, C.P., and Morrissey, J.P. (2011). Physiological and metabolic diversity in the yeast Kluyveromyces marxianus. Antonie van Leeuwenhoek 100, 507–519. 10.1007/s10482-011-9606-x.

6. Varela, J.A., Puricelli, M., Ortiz-Merino, R.A., Giacomobono, R., Braun-Galleani, S., Wolfe, K.H., and Morrissey, J.P. (2019). Origin of Lactose Fermentation in Kluyveromyces lactis by Interspecies Transfer of a Neo-functionalized Gene Cluster during Domestication. Curr Biol 29, 4284–4290 e4282. 10.1016/j.cub.2019.10.044.

7. Marcus, J.F., DeMarsh, T.A., and Alcaine, S.D. (2021). Upcycling of Whey Permeate through Yeast- and Mold-Driven Fermentations under Anoxic and Oxic Conditions. Fermentation 7, 16.

8. Nascimento, M.F., Barreiros, R., Oliveira, A.C., Ferreira, F.C., and Faria, N.T. (2022). Moesziomyces spp. cultivation using cheese whey: new yeast extract-free media, β-galactosidase biosynthesis and mannosylerythritol lipids production. Biomass Conversion and Biorefinery. 10.1007/s13399-022-02837-y.

9. Thoden, J.B., and Holden, H.M. (2007). The molecular architecture of glucose-1-phosphate uridylyltransferase. Protein Sci 16, 432–440. 10.1110/ps.062626007.

10. Slot, J.C., and Rokas, A. (2010). Multiple GAL pathway gene clusters evolved independently and by different mechanisms in fungi. Proc Natl Acad Sci U S A 107, 10136–10141. 10.1073/pnas.0914418107.

11. Mojzita, D., Herold, S., Metz, B., Seiboth, B., and Richard, P. (2012). l-xylo-3-Hexulose Reductase Is the Missing Link in the Oxidoreductive Pathway for d-Galactose Catabolism in Filamentous Fungi*. Journal of Biological Chemistry 287, 26010–26018. 10.1074/jbc.M112.372755.

12. 12. Gruben, B.S., Zhou, M., and de Vries, R.P. (2012). GalX regulates the D-galactose oxido-reductive pathway in Aspergillus niger. FEBS Lett 586, 3980–3985. 10.1016/j.febslet.2012.09.029.

13. Liu, J.J., Zhang, G.C., Kwak, S., Oh, E.J., Yun, E.J., Chomvong, K., Cate, J.H.D., and Jin, Y.S. (2019). Overcoming the thermodynamic equilibrium of an isomerization reaction through oxidoreductive reactions for biotransformation. Nat Commun 10, 1356. 10.1038/s41467-019-09288-6.

14. Zhang, G., Zabed, H.M., An, Y., Yun, J., Huang, J., Zhang, Y., Li, X., Wang, J., Ravikumar, Y., and Qi, X. (2022). Biocatalytic conversion of a lactose-rich dairy waste into D-tagatose, D-arabitol and galactitol using sequential whole cell and fermentation technologies. Bioresource Technology 358, 127422. 10.1016/j.biortech.2022.127422.

15. Jagtap, S.S., Bedekar, A.A., Liu, J.J., Jin, Y.S., and Rao, C.V. (2019). Production of galactitol from galactose by the oleaginous yeast Rhodosporidium toruloides IFO0880. Biotechnol Biofuels 12, 250. 10.1186/s13068-019-1586-5.

16. Harrison, M.C., LaBella, A.L., Hittinger, C.T., and Rokas, A. (2022). The evolution of the GALactose utilization pathway in budding yeasts. Trends Genet 38, 97–106. 10.1016/j.tig.2021.08.013.

17. Rokas, A., Wisecaver, J.H., and Lind, A.L. (2018). The birth, evolution and death of metabolic gene clusters in fungi. Nature Reviews Microbiology 16, 731–744. 10.1038/s41579-018-0075-3.

18. Wong, S., and Wolfe, K.H. (2005). Birth of a metabolic gene cluster in yeast by adaptive gene relocation. Nat Genet 37, 777–782. 10.1038/ng1584.

19. Krause, D.J., Kominek, J., Opulente, D.A., Shen, X.-X., Zhou, X., Langdon, Q.K., DeVirgilio, J., Hulfachor, A.B., Kurtzman, C.P., Rokas, A., and Hittinger, C.T. (2018). Functional and evolutionary characterization of a secondary metabolite gene cluster in budding yeasts. Proceedings of the National Academy of Sciences 115, 11030–11035. doi:10.1073/pnas.1806268115.

20. 20. de Jongh, W.A., Bro, C., Ostergaard, S., Regenberg, B., Olsson, L., and Nielsen, J. (2008). The roles of galactitol, galactose-1-phosphate, and phosphoglucomutase in galactose-induced toxicity in Saccharomyces cerevisiae. Biotechnol Bioeng 101, 317–326. 10.1002/bit.21890.

21. Lawrence, J. (1999). Selfish operons: the evolutionary impact of gene clustering in prokaryotes and eukaryotes. Current Opinion in Genetics & Development 9, 642–648. 10.1016/S0959-437X(99)00025-8.

22. Martchenko, M., Levitin, A., Hogues, H., Nantel, A., and Whiteway, M. (2007). Transcriptional rewiring of fungal galactose-metabolism circuitry. Curr Biol 17, 1007–1013. 10.1016/j.cub.2007.05.017.

23. 23. Van Ende, M., Wijnants, S., and Van Dijck, P. (2019). Sugar Sensing and Signaling in Candida albicans and Candida glabrata. Front Microbiol 10, 99. 10.3389/fmicb.2019.00099.

24. Peng, G., and Hopper, J.E. (2002). Gene activation by interaction of an inhibitor with a cytoplasmic signaling protein. Proceedings of the National Academy of Sciences 99, 8548–8553. 10.1073/pnas.142100099.

25. Bhat, P.J., and Murthy, T.V. (2001). Transcriptional control of the GAL/MEL regulon of yeast Saccharomyces cerevisiae: mechanism of galactose-mediated signal transduction. Mol Microbiol 40, 1059–1066. 10.1046/j.1365-2958.2001.02421.x.

26. Meyer, J., Walker-Jonah, A., and Hollenberg, C.P. (1991). Galactokinase encoded by GAL1 is a bifunctional protein required for induction of the GAL genes in Kluyveromyces lactis and is able to suppress the gal3 phenotype in Saccharomyces cerevisiae. Molecular and Cellular Biology 11, 5454–5461. doi:10.1128/mcb.11.11.5454-5461.1991.

27. Halvorsen, Y.C., Nandabalan, K., and Dickson, R.C. (1990). LAC9 DNA-binding domain coordinates two zinc atoms per monomer and contacts DNA as a dimer. Journal of Biological Chemistry 265, 13283–13289. 10.1016/S0021-9258(19)38296-1.

28. Dalal, C.K., Zuleta, I.A., Mitchell, K.F., Andes, D.R., El-Samad, H., and Johnson, A.D. (2016). Transcriptional rewiring over evolutionary timescales changes quantitative and qualitative properties of gene expression. Elife 5. 10.7554/eLife.18981.

29. Sun, X., Yu, J., Zhu, C., Mo, X., Sun, Q., Yang, D., Su, C., and Lu, Y. (2023). Recognition of galactose by a scaffold protein recruits a transcriptional activator for the GAL regulon induction in Candida albicans. Elife 12. 10.7554/eLife.84155.

30. Wu, J., Hu, J., Zhao, S., He, M., Hu, G., Ge, X., and Peng, N. (2018). Single-cell Protein and Xylitol Production by a Novel Yeast Strain *Candida intermedia* FL023 from Lignocellulosic Hydrolysates and Xylose. Appl Biochem Biotechnol 185, 163–178. 10.1007/s12010-017-2644-8.

31. Gardonyi, M., Osterberg, M., Rodrigues, C., Spencer-Martins, I., and Hahn-Hagerdal, B. (2003). High capacity xylose transport in Candida intermedia PYCC 4715. FEMS Yeast Res 3, 45–52. 10.1111/j.1567-1364.2003.tb00137.x.

32. Fonseca, C., Olofsson, K., Ferreira, C., Runquist, D., Fonseca, L.L., Hahn-Hagerdal, B., and Liden, G. (2011). The glucose/xylose facilitator Gxf1 from *Candida intermedia* expressed in a xylose-fermenting industrial strain of Saccharomyces cerevisiae increases xylose uptake in SSCF of wheat straw. Enzyme Microb Technol 48, 518–525. 10.1016/j.enzmictec.2011.02.010.

33. Moreno, A.D., Carbone, A., Pavone, R., Olsson, L., and Geijer, C. (2019). Evolutionary engineered Candida intermedia exhibits improved xylose utilization and robustness to lignocellulose-derived inhibitors and ethanol. Appl Microbiol Biotechnol 103, 1405–1416. 10.1007/s00253-018-9528-x.

34. Mayr, P., Bruggler, K., Kulbe, K.D., and Nidetzky, B. (2000). D-Xylose metabolism by *Candida intermedia*: isolation and characterisation of two forms of aldose reductase with different coenzyme specificities. J Chromatogr B Biomed Sci Appl 737, 195–202. 10.1016/s0378-4347(99)00380-1.

35. Nidetzky, B., Bruggler, K., Kratzer, R., and Mayr, P. (2003). Multiple forms of xylose reductase in *Candida intermedia*: comparison of their functional properties using quantitative structure-activity relationships, steady-state kinetic analysis, and pH studies. J Agric Food Chem 51, 7930–7935. 10.1021/jf034426j.

36. Yonten, V., and Aktas, N. (2014). Exploring the optimum conditions for maximizing the microbial growth of Candida intermedia by response surface methodology. Prep Biochem Biotechnol 44, 26–39. 10.1080/10826068.2013.782044.

37. Geijer, C., Faria-Oliveira, F., Moreno, A.D., Stenberg, S., Mazurkewich, S., and Olsson, L. (2020). Genomic and transcriptomic analysis of *Candida intermedia* reveals the genetic determinants for its xylose-converting capacity. Biotechnol Biofuels 13, 48. 10.1186/s13068-020-1663-9.

38. Moreno, A.D., Tellgren-Roth, C., Soler, L., Dainat, J., Olsson, L., and Geijer, C. (2017). Complete Genome Sequences of the Xylose-Fermenting Candida intermedia Strains CBS 141442 and PYCC 4715. Genome Announc 5. 10.1128/genomeA.00138-17.

39. Peri, K.V.R., Faria-Oliveira, F., Larsson, A., Plovie, A., Papon, N., and Geijer, C. (2023). Split-marker-mediated genome editing improves homologous recombination frequency in the CTG clade yeast Candida intermedia. FEMS Yeast Res 23. 10.1093/femsyr/foad016.

40. Wray, L.V., Jr., Witte, M.M., Dickson, R.C., and Riley, M.I. (1987). Characterization of a positive regulatory gene, LAC9, that controls induction of the lactose-galactose regulon of Kluyveromyces lactis: structural and functional relationships to GAL4 of Saccharomyces cerevisiae. Mol Cell Biol 7, 1111–1121. 10.1128/mcb.7.3.1111-1121.1987.

41. Douglas, H.C., and Hawthorne, D.C. (1964). ENZYMATIC EXPRESSION AND GENETIC LINKAGE OF GENES CONTROLLING GALACTOSE UTILIZATION IN SACCHAROMYCES. Genetics 49, 837–844. 10.1093/genetics/49.5.837.

42. Bailey, T.L., Johnson, J., Grant, C.E., and Noble, W.S. (2015). The MEME Suite. Nucleic Acids Research 43, W39–W49. 10.1093/nar/gkv416.

43. Thoden, J.B., Sellick, C.A., Timson, D.J., Reece, R.J., and Holden, H.M. (2005). Molecular structure of Saccharomyces cerevisiae Gal1p, a bifunctional galactokinase and transcriptional inducer. J Biol Chem 280, 36905–36911. 10.1074/jbc.M508446200.

44. Jumper, J., Evans, R., Pritzel, A., Green, T., Figurnov, M., Ronneberger, O., Tunyasuvunakool, K., Bates, R., Zidek, A., Potapenko, A., et al. (2021). Highly accurate protein structure prediction with AlphaFold. Nature 596, 583–589. 10.1038/s41586-021-03819-2.

45. Hittinger, C.T., and Carroll, S.B. (2007). Gene duplication and the adaptive evolution of a classic genetic switch. Nature 449, 677–681. 10.1038/nature06151.

46. Geronikou, A., Larsen, N., Lillevang, S.K., and Jespersen, L. (2022). Occurrence and Identification of Yeasts in Production of White-Brined Cheese. Microorganisms 10, 1079.

47. Tanji, M., Namimatsu, K., Kinoshita, M., Motoshima, H., Oda, Y., and Ohnishi, M. (2004). Content and Chemical Compositions of Cerebrosides in Lactose-assimilating Yeasts. Bioscience, Biotechnology, and Biochemistry 68, 2205–2208. 10.1271/bbb.68.2205.

48. Yano, K., and Fukasawa, T. (1997). Galactose-dependent reversible interaction of Gal3p with Gal80p in the induction pathway of Gal4p-activated genes of Saccharomyces cerevisiae. Proc Natl Acad Sci U S A 94, 1721–1726. 10.1073/pnas.94.5.1721.

49. Tesfaw, A., Oner, E.T., and Assefa, F. (2021). Evaluating crude whey for bioethanol production using non-Saccharomyces yeast, Kluyveromyces marxianus. SN Applied Sciences 3, 42. 10.1007/s42452-020-03996-1.

50. Maullu, C., Lampis, G., Desogus, A., Ingianni, A., Rossolini, G.M., and Pompei, R. (1999). High-level production of heterologous protein by engineered yeasts grown in cottage cheese whey. Appl Environ Microbiol 65, 2745–2747. 10.1128/aem.65.6.2745-2747.1999.

51. Urit, T., Stukert, A., Bley, T., and Löser, C. (2012). Formation of ethyl acetate by Kluyveromyces marxianus on whey during aerobic batch cultivation at specific trace element limitation. Appl Microbiol Biotechnol 96, 1313–1323. 10.1007/s00253-012-4107-z.

52. Santana, D.J., and O’Meara, T.R. (2021). Forward and reverse genetic dissection of morphogenesis identifies filament-competent Candida auris strains. Nat Commun 12, 7197. 10.1038/s41467-021-27545-5.

53. Gietz, R.D., and Schiestl, R.H. (2007). High-efficiency yeast transformation using the LiAc/SS carrier DNA/PEG method. Nature Protocols 2, 31–34. 10.1038/nprot.2007.13.

54. Verduyn, C., Postma, E., Scheffers, W.A., and Van Dijken, J.P. (1992). Effect of benzoic acid on metabolic fluxes in yeasts: a continuous-culture study on the regulation of respiration and alcoholic fermentation. Yeast 8, 501–517. 10.1002/yea.320080703.

55. Bruder, S., Reifenrath, M., Thomik, T., Boles, E., and Herzog, K. (2016). Parallelised online biomass monitoring in shake flasks enables efficient strain and carbon source dependent growth characterisation of Saccharomyces cerevisiae. Microb Cell Fact 15, 127. 10.1186/s12934-016-0526-3.

56. Bruder, S., Reifenrath, M., Thomik, T., Boles, E., and Herzog, K. (2016). Parallelised online biomass monitoring in shake flasks enables efficient strain and carbon source dependent growth characterisation of Saccharomyces cerevisiae. Microbial Cell Factories 15, 127. 10.1186/s12934-016-0526-3.

57. Goncalves, C., Wisecaver, J.H., Kominek, J., Oom, M.S., Leandro, M.J., Shen, X.X., Opulente, D.A., Zhou, X., Peris, D., Kurtzman, C.P., et al. (2018). Evidence for loss and reacquisition of alcoholic fermentation in a fructophilic yeast lineage. Elife 7. 10.7554/eLife.33034.

58. Katoh, K., and Standley, D.M. (2013). MAFFT multiple sequence alignment software version 7: improvements in performance and usability. Mol Biol Evol 30, 772–780. 10.1093/molbev/mst010.

59. Capella-Gutiérrez, S., Silla-Martínez, J.M., and Gabaldón, T. (2009). trimAl: a tool for automated alignment trimming in large-scale phylogenetic analyses. Bioinformatics 25, 1972–1973. 10.1093/bioinformatics/btp348.

60. 60. Nguyen, L.T., Schmidt, H.A., von Haeseler, A., and Minh, B.Q. (2015). IQ-TREE: a fast and effective stochastic algorithm for estimating maximum-likelihood phylogenies. Mol Biol Evol 32, 268–274. 10.1093/molbev/msu300.

61. 61. Minh, B.Q., Nguyen, M.A., and von Haeseler, A. (2013). Ultrafast approximation for phylogenetic bootstrap. Mol Biol Evol 30, 1188–1195. 10.1093/molbev/mst024.

62. Letunic, I., and Bork, P. (2019). Interactive Tree Of Life (iTOL) v4: recent updates and new developments. Nucleic Acids Res 47, W256–w259. 10.1093/nar/gkz239.

63. Gupta, S., Stamatoyannopoulos, J.A., Bailey, T.L., and Noble, W.S. (2007). Quantifying similarity between motifs. Genome Biology 8, R24. 10.1186/gb-2007-8-2-r24.

64. Teixeira, M.C., Viana, R., Palma, M., Oliveira, J., Galocha, M., Mota, M.N., Couceiro, D., Pereira, M.G., Antunes, M., Costa, I.V., et al. (2022). YEASTRACT+: a portal for the exploitation of global transcription regulation and metabolic model data in yeast biotechnology and pathogenesis. Nucleic Acids Research 51, D785–D791. 10.1093/nar/gkac1041.

65. Andrews, S. (2010). FastQC: a quality control tool for high throughput sequence data. Babraham Bioinformatics, Babraham Institute, Cambridge, United Kingdom.

66. Dobin, A., Davis, C.A., Schlesinger, F., Drenkow, J., Zaleski, C., Jha, S., Batut, P., Chaisson, M., and Gingeras, T.R. (2013). STAR: ultrafast universal RNA-seq aligner. Bioinformatics 29, 15–21.

67. Robinson, M.D., McCarthy, D.J., and Smyth, G.K. (2010). edgeR: a Bioconductor package for differential expression analysis of digital gene expression data. bioinformatics 26, 139–140.

68. Smyth, G.K. (2005). Limma: linear models for microarray data. In Bioinformatics and computational biology solutions using R and Bioconductor, (Springer), pp. 397–420.

69. Love, M.I., Huber, W., and Anders, S. (2014). Moderated estimation of fold change and dispersion for RNA-seq data with DESeq2. Genome biology 15, 1–21.

